# Regulatory T cell activation triggers specific changes in glycosylation associated with Siglec-1-dependent inflammatory responses

**DOI:** 10.1101/2020.07.29.226399

**Authors:** Gang Wu, Gavuthami Murugesan, Manjula Nagala, Alex McCraw, Stuart M. Haslam, Anne Dell, Paul R. Crocker

**Author notes:** Department of Life Sciences, Imperial College London, London, United Kingdom (Stuart M. Haslam, Anne Dell).

## Abstract

Siglec-1 is a macrophage lectin-like receptor that mediates sialic acid-dependent cellular interactions. It was shown previously to promote inflammation in autoimmune disease through suppressing the expansion of regulatory T cells (Tregs). We have investigated the molecular basis for Siglec-1 binding to these cells using *in vitro*-induced Tregs. Siglec-1 binding was strongly upregulated on activated cells, but lost under resting conditions. Glycosylation changes that affect Siglec-1 binding were studied by comparing activated and resting Tregs using RNA-Seq, glycomics, proteomics and binding of selected antibodies and lectins. A proximity labelling and proteomics strategy identified 49 glycoproteins expressed by activated Tregs that may function as Siglec-1 counter-receptors. These represent ∼5% of the total membrane protein pool and were mainly related to T cell activation and proliferation. We demonstrate that several of these counter-receptors are upregulated following activation of Tregs and provide initial evidence that their altered glycosylation may also be important for Siglec-1 binding.

## Introduction

All mammalian cells are coated with a dense layer of glycans termed the glycocalyx (Tarbell & Cancel, 2016). Despite the diverse structures and inherent complexity of these glycans, they are frequently capped with sialic acid moieties. Sialic acids can mediate a wide variety of functions (Varki, Schnaar, & Schauer, 2017), but an important feature is that they serve as ligands for a family of endogenous sialic acid binding lectins of the Ig superfamily known as Siglecs (Crocker, Paulson, & Varki, 2007). The interaction between Siglecs and their ligands can regulate the functional activities of most cells of the immune system (Crocker et al., 2007; Macauley, Crocker, & Paulson, 2014). Siglec-1 (also known as sialoadhesin, Sn or CD169) is a macrophage-restricted Siglec that is well conserved across mammals (Klaas & Crocker, 2012). Under normal physiological conditions, it is highly expressed on macrophage subsets in secondary lymphoid tissues and its expression on other macrophages can be induced at sites of inflammation (Klaas & Crocker, 2012; Schadee-Eestermans et al., 2000). Siglec-1 appears to have evolved primarily as a cellular interaction molecule. It has an unusually large number of 17 Ig domains that are thought to project the sialic acid binding site away from the plasma membrane to promote interactions with sialic acid ligands presented on other cells (Klaas & Crocker, 2012). This is in striking contrast to other Siglecs which have between 2 and 7 Ig domains and are typically masked at the cell surface by interactions with sialic acids in *cis* (Crocker et al., 2007).

In addition to being displayed on cells of the host, the sialic acids recognised by Siglec-1 can be present on certain pathogens such as sialylated bacteria, protozoa and enveloped viruses and their recognition can lead to increased pathogen uptake by macrophages and enhanced host susceptibility (reviewed in (Macauley et al., 2014)). However, the predominant biological functions of Siglec-1 involve interactions with sialic acids of the host. For example, Siglec-1 can mediate sialic acid-dependent crosstalk between macrophages and various immune cells including neutrophils (Crocker, Freeman, Gordon, & Kelm, 1995), dendritic cells (van Dinther et al., 2018), innate-like lymphocytes (Zhang et al., 2016) and regulatory T cells (C. Wu et al., 2009).

An important biological function of Siglec-1, discovered in studies of Siglec-1-deficient mice, is its role in promoting inflammatory responses during various autoimmune diseases of the nervous system (Groh et al., 2016; Ip, Kroner, Crocker, Nave, & Martini, 2007; Jiang et al., 2006; Kobsar et al., 2006; C. Wu et al., 2009). Mechanistically, this is likely to be due to Siglec-1-dependent suppression of Treg expansion. This was initially implied in studies of experimental autoimmune uveitis, which showed that Siglec-1 was a prominent marker on inflammatory macrophages at the peak phase of tissue damage (Jiang et al., 2006). Siglec-1-deficient mice exhibited reduced disease severity and decreased proliferation and IFN-γ secretion by effector T cells. Direct evidence for an important cross-talk between Siglec-1 and Tregs was seen in a study of experimental autoimmune encephalomyelitis (EAE), which is a mouse model of multiple sclerosis (C. Wu et al., 2009). The EAE model revealed that Siglec-1-expressing macrophages are closely associated with activated CD4+Foxp3+ Tregs at sites of inflammation within the central nervous system. Siglec-1-deficient mice had increased numbers of Tregs and reduced levels of Th17 cells producing inflammatory cytokines, leading to attenuated disease severity (C. Wu et al., 2009). The Tregs isolated from diseased mice showed strong sialic acid-dependent binding to Siglec-1 and co-culture with macrophages suppressed their expansion in a Siglec-1-dependent manner. Similar results have been observed in a mouse model of neuronal ceroid lipofuscinoses, which showed that Siglec-1 negatively controls CD8+CD122+ regulatory T cells, and promotes neuroinflammation-related disease progression (Groh et al., 2016).

A major gap in our understanding of how Siglecs modulate cellular functions in the immune system is the identification of endogenous ligands and counter-receptors on relevant cell populations. Here, the ligand is defined as the oligosaccharide structure recognised by Siglec-1 and the counter-receptor is the composite of the ligand(s) attached to an appropriate protein or lipid carrier (Crocker & Feizi, 1996). Using defined glycans, Siglec-1 has been found to prefer α2,3-linked Neu5Ac over α2,6- and α2,8-linked Neu5Ac (Crocker et al., 1991; Crocker, Vinson, Kelm, & Drickamer, 1999). Certain other types of sialic acid, including Neu5Gc and Neu5,9(Ac)_2_, were not recognised by Siglec-1 (Kelm, Schauer, Manuguerra, Gross, & Crocker, 1994). Like many membrane lectins, Siglec-1 exhibits low binding affinities for its glycan ligands, with Kd values in the millimolar range (Crocker et al., 1999). Cell adhesion mediated by Siglec-1 therefore depends on the clustering of both Siglec-1 and its ligands on cell surfaces to obtain high avidity interactions. As a result, the molecular basis for Siglec-1 binding to Tregs is complex and determined by multiple factors. On the one hand it can be affected by global factors involved in the synthesis of glycan ligands, such as the production and transport of sugar donors and the expression of sialyltransferases and other glycosyl transferases. On the other hand, it can also be affected by specific factors, such as the expression and localisation of particular glycoprotein and glycolipid counter-receptors which carry the glycan ligands.

The aim of this study was to investigate the global and specific factors that lead to Siglec-1 binding to activated Tregs using RNA-Seq, glycomics and proteomics. A proximity labeling strategy, combined with proteomics, was used to identify glycoprotein counter-receptors for Siglec-1 expressed by activated Tregs.

## Results

### Siglec-1 binding to Tregs depends completely on Treg activation

FoxP3-positive CD4 Tregs are a small subset of the total pool of CD4 T cells but can be induced from FoxP3-negative CD4 T cells under defined culture conditions. In order to obtain a sufficient number of Tregs for RNA-Seq, glycomics and proteomics, CD4 T cells were isolated from mouse spleens and lymph nodes and induced to become Tregs as illustrated in Figure 1. After induction for 4 – 5 days, the proportion of Tregs increased from about 4% to about 90%, and the cell count increased 1- to 3-fold. After expansion for a further 4 days, the cell count increased another 2- to 3-fold, without affecting the proportion of Tregs (Figure 1). Siglec-1 binding to these Tregs was analysed by flow cytometry. The freshly-induced and activated Tregs exhibited strong Siglec-1 binding (Figure 2A). When these cells were cultured under resting conditions in the absence of anti-CD3 antibody, Siglec-1 binding disappeared completely, but was fully restored when the cells were re-stimulated with anti-CD3 antibody (Figure 2A). Alternatively, when the freshly-induced Tregs were kept activated with anti-CD3, they continued to exhibit strong Siglec-1 binding until anti-CD3 was removed from the cell culture medium (Figure 2A). The extent of induction of Siglec-1 ligands on Tregs by anti-CD3 antibody and its kinetics were dose-dependent over a range of anti-CD3 concentrations (Figure 2B). These observations reveal that Siglec-1 binding to Tregs *in vitro* depends completely on Treg activation. This is consistent with previous *in vivo* studies showing that Siglec-1 bound only to activated, but not resting Tregs (Kidder, Richards, Ziltener, Garden, & Crocker, 2013; C. Wu et al., 2009).

**Figure 1.**
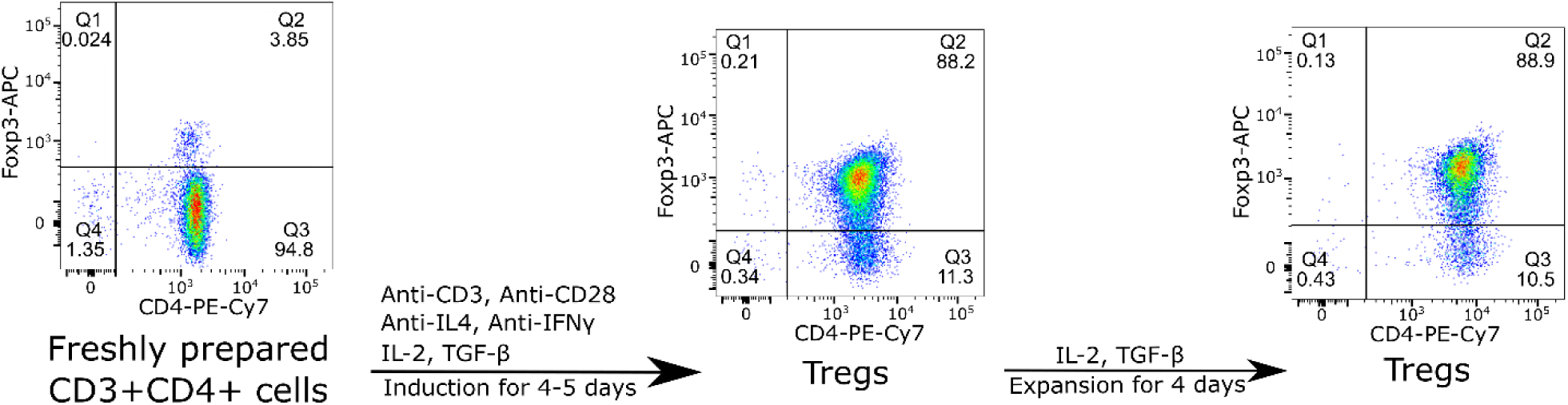
Flow cytometry showing induction of FoxP3+ CD4+Tregs and their expansion *in vitro*. Mouse CD4 T cells were isolated from spleen and lymph nodes, stimulated with anti-CD3 and anti-CD28 mAbs for 4-5 days in the presence of anti-IL-4, anti-IFNγ, IL-2 and TGF-β. The cells were then expanded in the presence of IL-2 and TGF-β for another 4 days. The data shown are representative of more than 10 experiments carried out.

**Figure 2.**
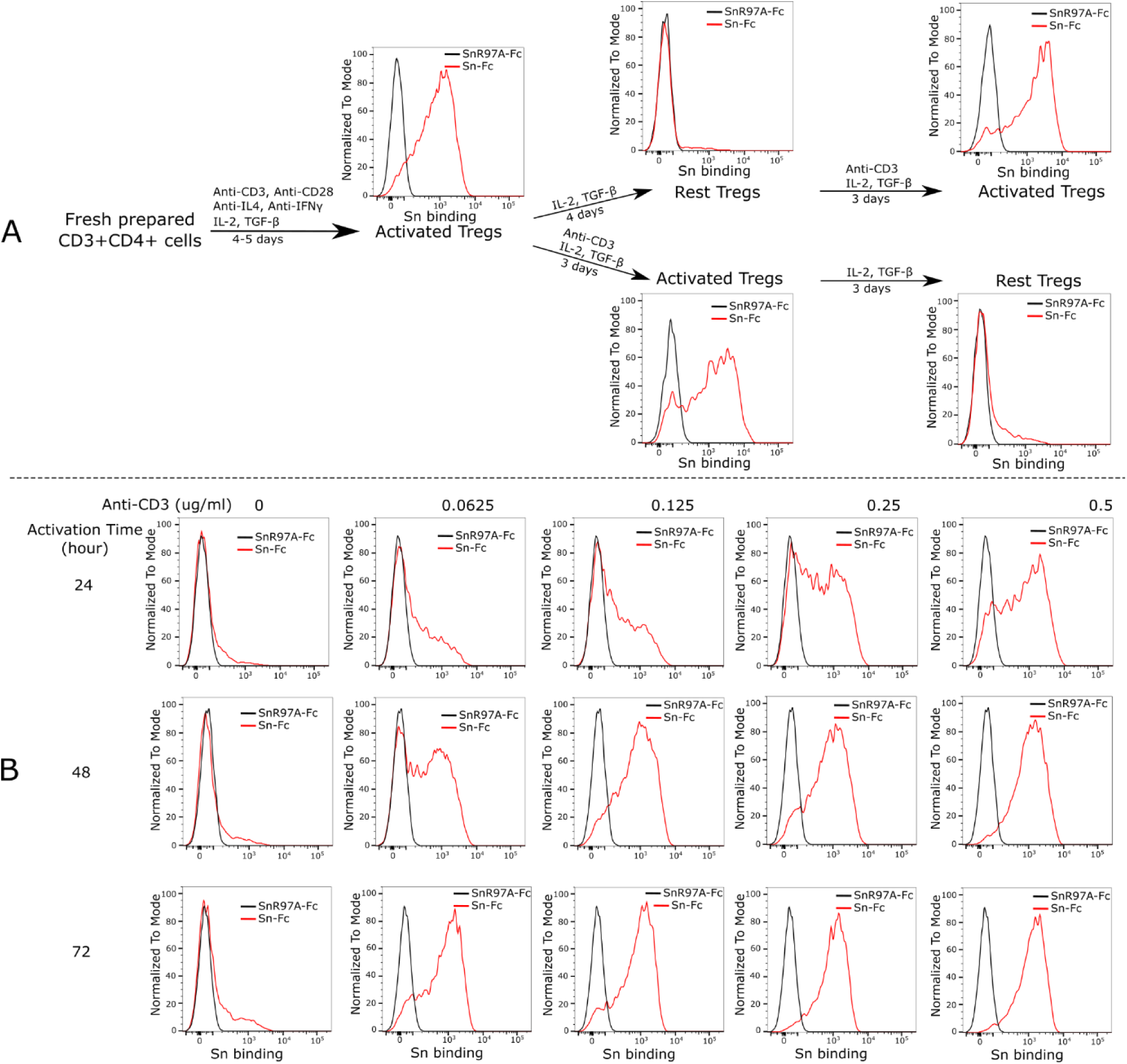
Siglec-1 binding to Tregs depends on Treg activation status. (**A)** Freshly isolated CD4 T cells were induced to become Tregs and analysed for expression of Siglec-1 ligands using pre-complexed Siglec-1-Fc (Sn-Fc). Compared to the negative control non-binding mutant, Siglec-1-R97A-Fc (SnR97A-Fc), the induced and activated Tregs showed strong Siglec-1 binding. The binding was lost when the cells were rested in IL-2 and TGF-β for 4 days, but was fully recovered when the cells were reactivated with anti-CD3 mAb for 3 days (upper panels). The freshly induced Tregs continued to show strong Siglec-1 binding when kept under activating conditions in the presence of anti-CD3 mAb, but binding was lost when anti-CD3 mAb was withdrawn for 3 days (lower panels). The whole set of experiments was performed twice with similar results. Analysis of Siglec-1 binding to resting and activated Tregs was repeated more than 20 times and in each case binding was much higher to the activated cells. (**B)** Siglec-1 binding to Tregs depends on the T cell receptor signal strength. Freshly induced Tregs were rested for 4 days and then stimulated using different concentrations of anti-CD3 mAb and analysed for Siglec-1 binding at 24, 48 and 72 hours. Binding was both dose- and time-dependent. Similar results were observed in 2 independent experiments.

### Siglec-1 binding to activated Tregs is not due to a global change of sialylation

The striking induction of Siglec-1 ligands on activated Tregs suggests that activation could be accompanied by global changes in glycan sialylation, as observed previously with CD4+ effector T cells (Naito-Matsui et al., 2014; Redelinghuys et al., 2011). These include increased expression of α2,3-linked sialic acids and a switch from the NeuGc to NeuAc form of sialic acid, both changes leading to increased binding of Siglec-1. To investigate this, we used plant lectins MAL II and SNA to probe the overall α2,3-linked and α2,6-linked sialylated glycans respectively, and anti-NeuGc antibody to measure NeuGc levels on Tregs. However, unlike effector T cells, there were no obvious changes in either lectin or antibody binding upon Treg activation (Figure 3).

**Figure 3.**
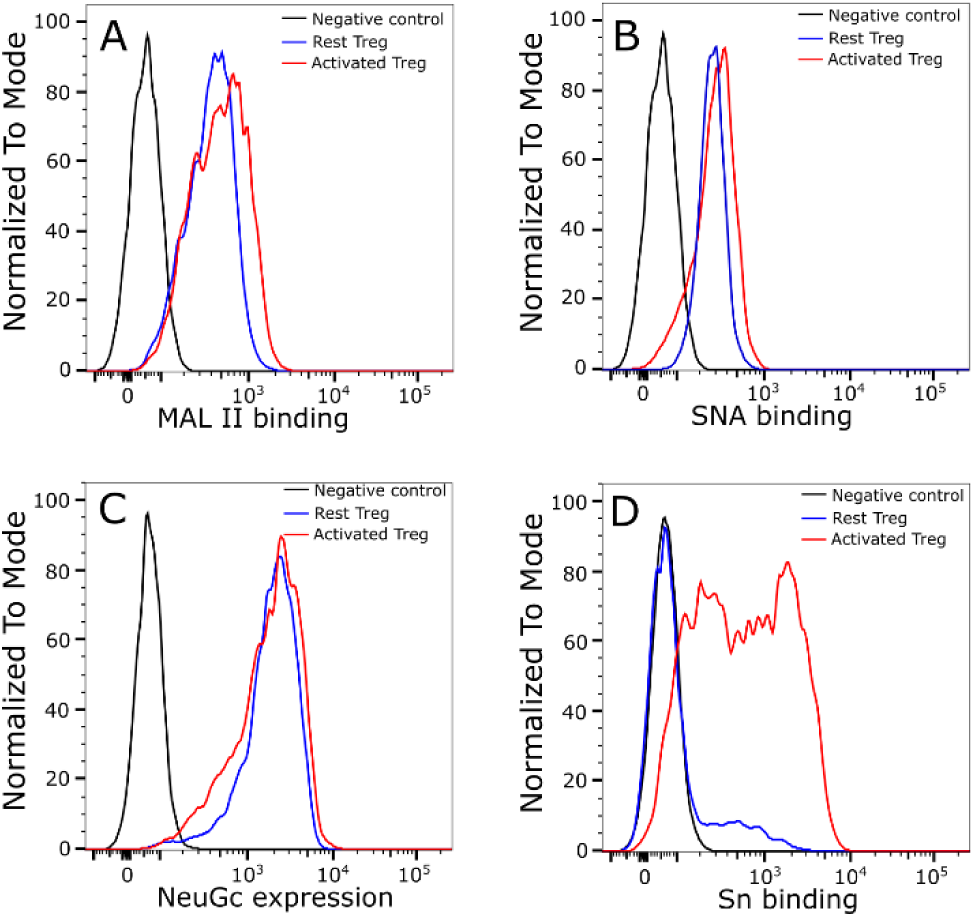
Treg activation does not lead to a global change of cell surface sialylation. α2,3-linked **(A)** and α2,6-linked sialic **(B)** acids on Tregs were probed using biotinylated plant lectins MAL II **(A)** or SNA **(B)**, which were then stained using FITC conjugated streptavidin for flow cytometry analysis. Tregs stained only with FITC conjugated streptavidin were used as a negative control. (**C)** NeuGc expression was analysed by flow cytometry using chicken IgY anti-NeuGc antibody. A chicken IgY isotype control antibody was used as a negative control. (**D)** Expression of Siglec-1 ligands was measured using complexes of Siglec-1-huIgG-Fc chimera mixed with FITC-conjugated goat anti-huIgG-Fc. Siglec-1-R97A-huIgG-Fc was used as a negative control. The same batch of Tregs was used for measurement of NeuGc and Siglec-1 ligand expression. The MAL II, SNA and anti-NeuGc experiments were performed twice and similar results were observed. Experiments to measure Siglec-1 binding to resting and activated Tregs were repeated more than 20 times and similar results were observed consistently.

Glycomics was used to directly examine the structures and relative quantities of Treg N-glycans and glycolipids. N-glycans showed an overall similar glycosylation and sialylation profile when resting and activated Tregs were compared (Figure 4). A specific change was found at m/z 3026, which is a bi-antennary glycan with core fucose and two NeuGc sialic acids. Relative to the other glycans, this glycan had a decreased intensity when Tregs became activated. Gal-α-Gal terminal structures were specifically found on mono-sialylated core fucosylated glycans at m/z 2809 and m/z 2839. Relative to the other biantennary glycans, the two glycans had a minor increase upon Treg activation. For glycolipids, MS (Figure 5) and MS/MS (Supplementary Figures 1-8) analyses showed different glycan profiles in resting and activated Tregs, with a trend of NeuGc switching to NeuAc upon activation. This can be seen by the higher proportions of NeuAc-containing glycans at m/z 1288, 1533 and 1649 versus their NeuGc-containing counterparts at m/z 1318, 1563 and 1709 in activated Tregs (Figure 5). This was especially striking for the α2,3-sialylated GM1b structures at m/z 1288 (NeuAc) and m/z 1318 (NeuGc) (Figure 5). These observations suggest that GM1b (NeuAc) is upregulated on activated Tregs where it could potentially serve as a ligand for Siglec-1.

**Figure 4.**
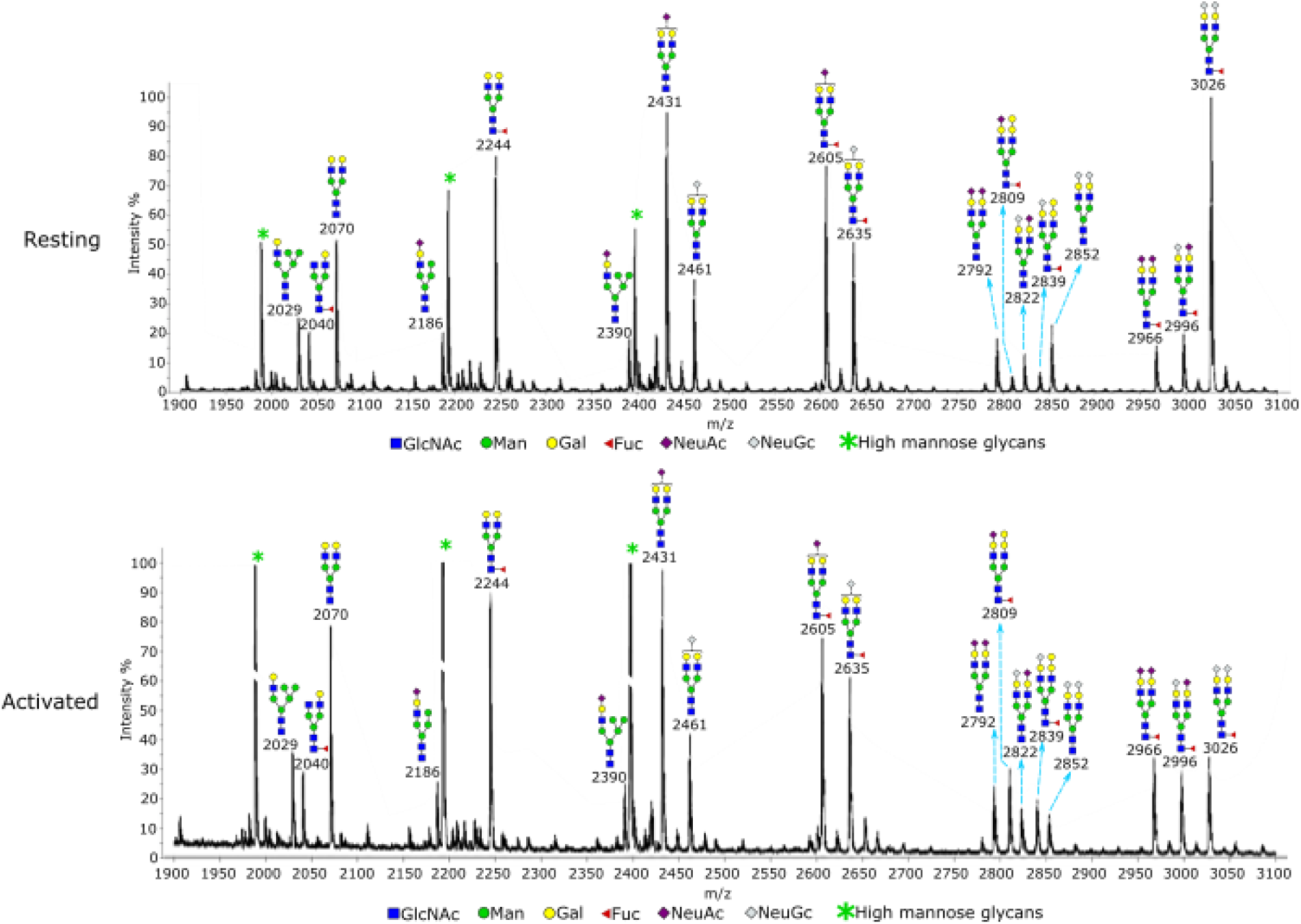
Glycomic analysis of resting and activated Tregs. N-glycans from the Tregs were permethylated and analysed by MALDI-TOF mass spectrometry. The data were acquired in the form of [M+Na]+ ions. Peaks representing hybrid and complex glycans were annotated according to the molecular weight and N-glycan biosynthetic pathways. Treg activation did not result in an overall change of glycosylation, except the glycan at m/z 3026, which had a decreased intensity relative to other glycans following activation of Tregs. A minor increase in Gal-α-Gal structure was observed upon Treg activation at m/z 2809 and m/z 2839, relative to the other biantennary glycans.

**Figure 5.**
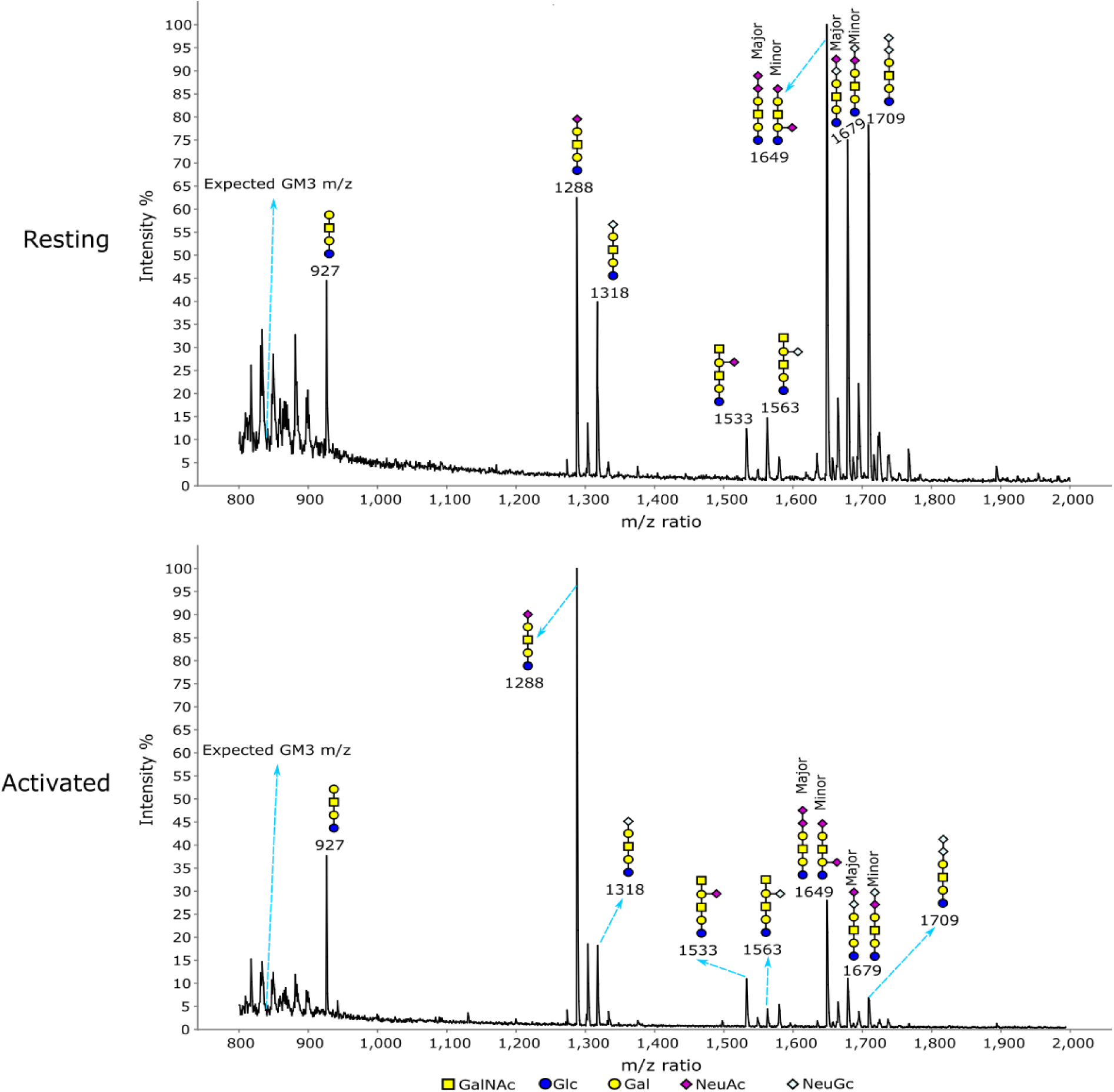
Glycomic analysis of glycolipid glycans on resting and activated Tregs. The data were annotated according to the molecular weight, biosynthetic pathways and MS/MS analysis. The NeuAc capped GM1b (m/z 1288) showed a much higher signal relative to other glycans in activated Tregs compared to resting Tregs.

### RNA-Seq analysis does not indicate a global change of sialylation in activated Tregs

RNA-Seq was used to profile the gene expression patterns, comparing resting and activated Tregs prepared from 4 independent biological replicates. The peak expression of Siglec-1 ligands was observed between 24 and 48 hours following Treg activation (Figure 2). Therefore we selected a 36 hour time point to isolate mRNA in order to maximise the chances of seeing clear changes in gene expression relevant to Siglec-1 ligand expression. The Pearson’s correlation coefficient for each pair of replicates is shown in the distance matrix in Figure 6A, which shows good quality data with reproducible replicates and major changes between resting and activated Tregs. This is further illustrated by clustering and PCA analyses (Figure 6B,C). To focus on genes involved in glycosylation, we assembled a dataset of 263 genes including glycosyltransferases, glycosidases, enzymes involved in amino sugar and nucleotide sugar metabolism, and sugar transporters (Supplementary Table 1). Whilst, overall, these genes did not show a dramatic global log2 fold change there were many specific changes upon Treg activation (Figure 6D, Supplementary Table 1). Mapping of the data to KEGG pathways points to overall increased N-glycosylation in activated Tregs (Supplementary Figure 9) and increased O-glycosylation for proteins that depend on expression of *Galnt3*, a GalNAc transferase that initiates O-glycosylation on serine and threonine residues (Supplementary Figure 10).

**Figure 6.**
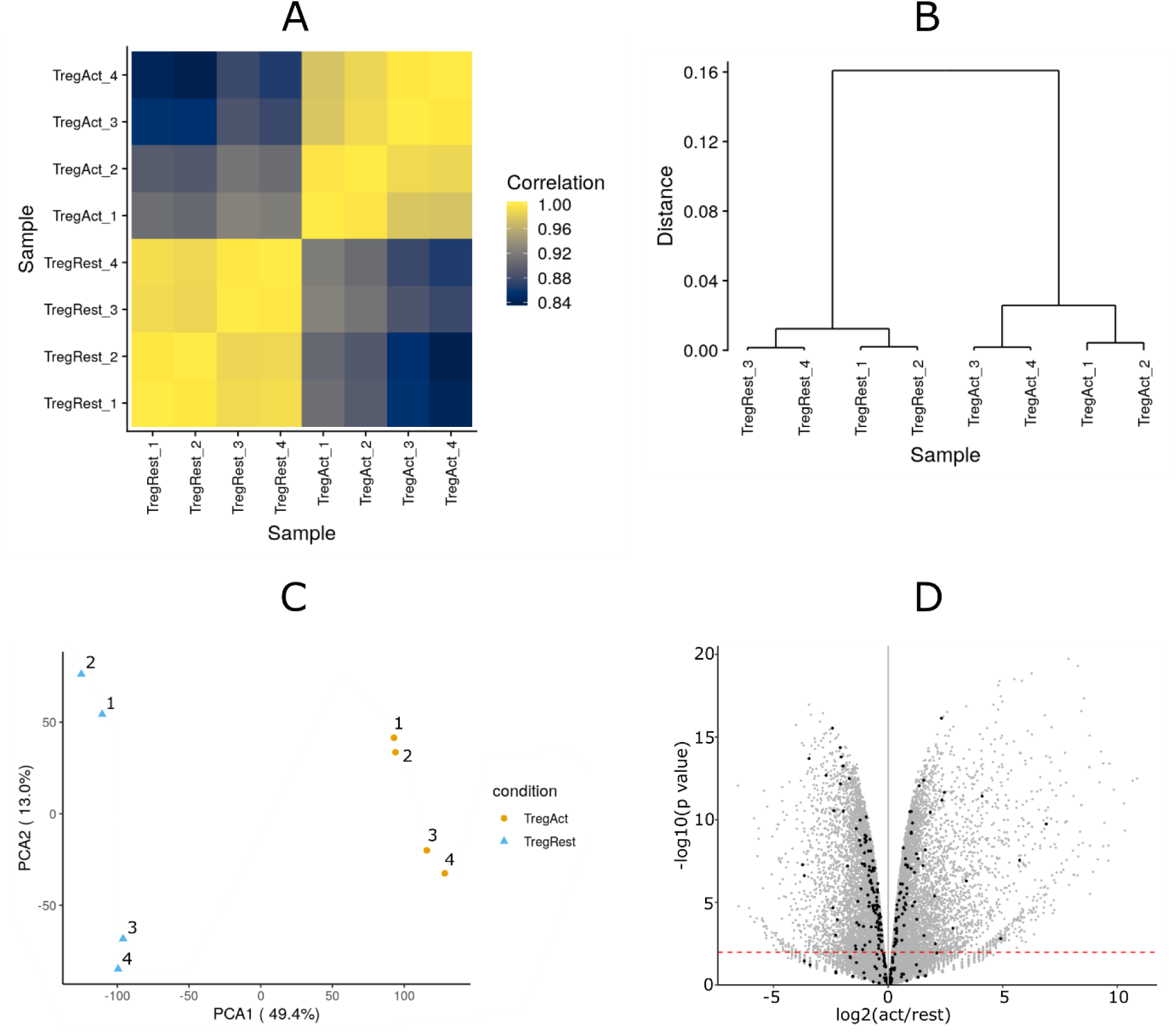
Overview of RNA-Seq data. Four biological replicates of resting and activated Tregs were prepared for RNA-Seq analysis. (**A-C)** Quality check of the RNA-Seq data shows high data quality with reproducible replicates and major changes between resting and activated Tregs. (**D)** Volcano plot of the RNA-Seq results. The black dots show glycosylation-related genes, while the grey dots show all other genes.

The expression patterns of genes involved in Treg sialylation are shown in Figure 7. The sialyltransferases (STs) are a family of ∼20 enzymes that transfer sialic acids to acceptor sugars in α2,3-, α2,6- and α2,8-glycosidic linkages (Takashima, 2008). For α2,3-sialylation, the preferred linkage for Siglec-1, Treg activation was associated with decreased expression of *St3gal1, St3gal2* and *St3gal6*, increased expression of *St3gal5* and no change for *St3gal3* and *St3gal4* (Figure 7A). Similarly, for α2,6-sialylation, differential expression was observed, with a 5-fold decrease of *St6gal1* and a 3-fold increase in expression of both *St6galnac4* and *St6galnac6* following activation (Figure 7B). The net effect of these alterations on overall α2,3 and α2,6 sialylation appears to be minimal as MALII and SNA staining for these respective linkages showed no changes on Treg activation, as described above (Figure 3A,B). For α2,8-sialylation, *St8sia1* and *St8sia4* were both decreased upon Treg activation, but expression of *St8sia6* was slightly increased (Figure 7C). We also analysed genes involved in modifications of sialic acids including conversion of NeuAc to 9-O-acetyl NeuAc and Neu5Gc, neither of which is recognised by Siglec-1 (Kelm et al., 1994). 9-O-acetylation is controlled by Casd1 and Siae which catalyse the addition or removal of acetyl groups to NeuAc respectively (Baumann et al., 2015; Orizio et al., 2015). Expression of both genes was reduced following Treg activation, suggesting no net change in levels of 9-O-acetyl NeuAc. (Figure 7D). Expression of *Cmah*, which is responsible for converting CMP-NeuAc to CMP-NeuGc (Kawano et al., 1995), did not change significantly upon Treg activation. This is consistent with the anti-NeuGc Ab binding results shown above (Figure 3C) which showed no differences between resting and activated Tregs. Finally, we analysed expression of endogenous sialidases with the potential to remove sialic acids from the cell surface. Of the 4 sialidase genes (Monti et al., 2010), only *Neu1* and *Neu3* were expressed in Tregs. Expression of *Neu1* increased slightly in activated Tregs while *Neu3* did not change significantly (Figure 7D). In conclusion, the results of RNA-Seq revealed changes in expression of many genes that affect glycosylation of multiple proteins and lipids, but did not reveal specific changes predicted to have major effects on Siglec-1 recognition following Treg activation.

**Figure 7.**
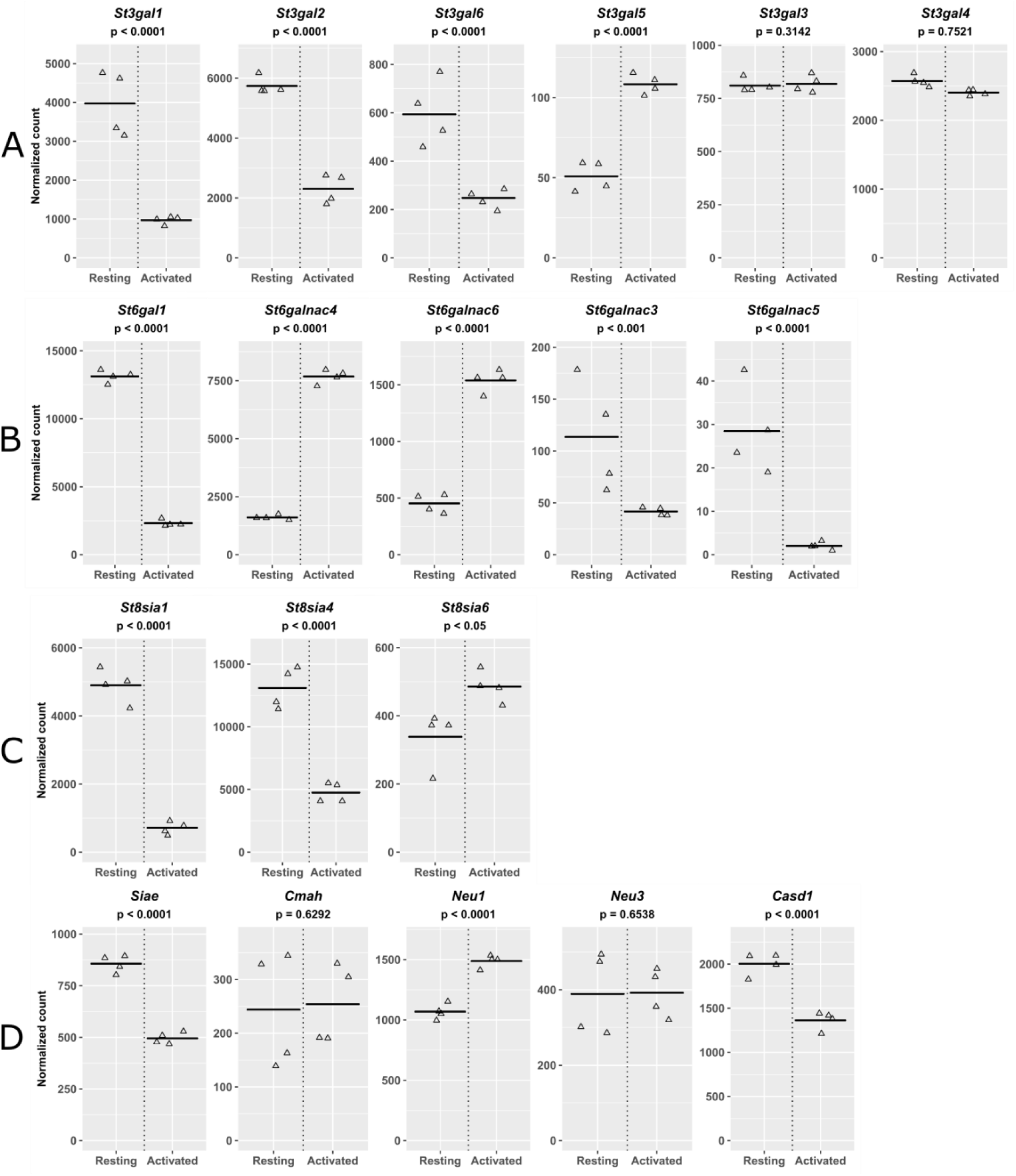
Normalized mRNA counts for genes involved in sialylation. Resting and activated Tregs from 4 biological replicates were analzyed by RNA-Seq. Genes involved in synthesis of **(A)** α2,3-linked sialic acid, **(B)** α2,6-linked sialic acid, **(C)** α2,8-linked sialic acid and **(D)** modification or removal of sialic acid are listed.

### Identification of glycoprotein counter-receptors for Siglec-1

A recently-described proximity labelling strategy (Chang et al., 2017; G. Wu, Nagala, & Crocker, 2017), coupled with quantitative proteomics, was used to identify Treg glycoproteins that could interact with Siglec-1. The experimental design is illustrated in Figure 8. It essentially involves preparation of Siglec-1-horse radish peroxidase (Sn-HRP) multimers (Figure 8A) that can bind to Tregs and, in the presence of tyramide-SS-biotin and H_2_O_2_, generate short-range biotin radicals that label neighbouring proteins (Figure 8B). Labelled proteins are then enriched with streptavidin and identified by quantitative proteomics (Figure 8C). To increase the chances of identifying diverse counter-receptors, two types of Siglec-1-HRP multimer were prepared, either in-solution, or attached to 50 nm microbeads (Figure 8A). Because of the potential for non-specific labelling and streptavidin binding, it was important to include similarly-prepared multimers of a non-binding negative control Siglec-1, (SnR97A) alongside the active Siglec-1. Proteins that were selectively enriched using the Sn-HRP complexes over the SnR97A-HRP complexes represent potential Siglec-1 counter-receptors (Figure 8C).

**Figure 8.**
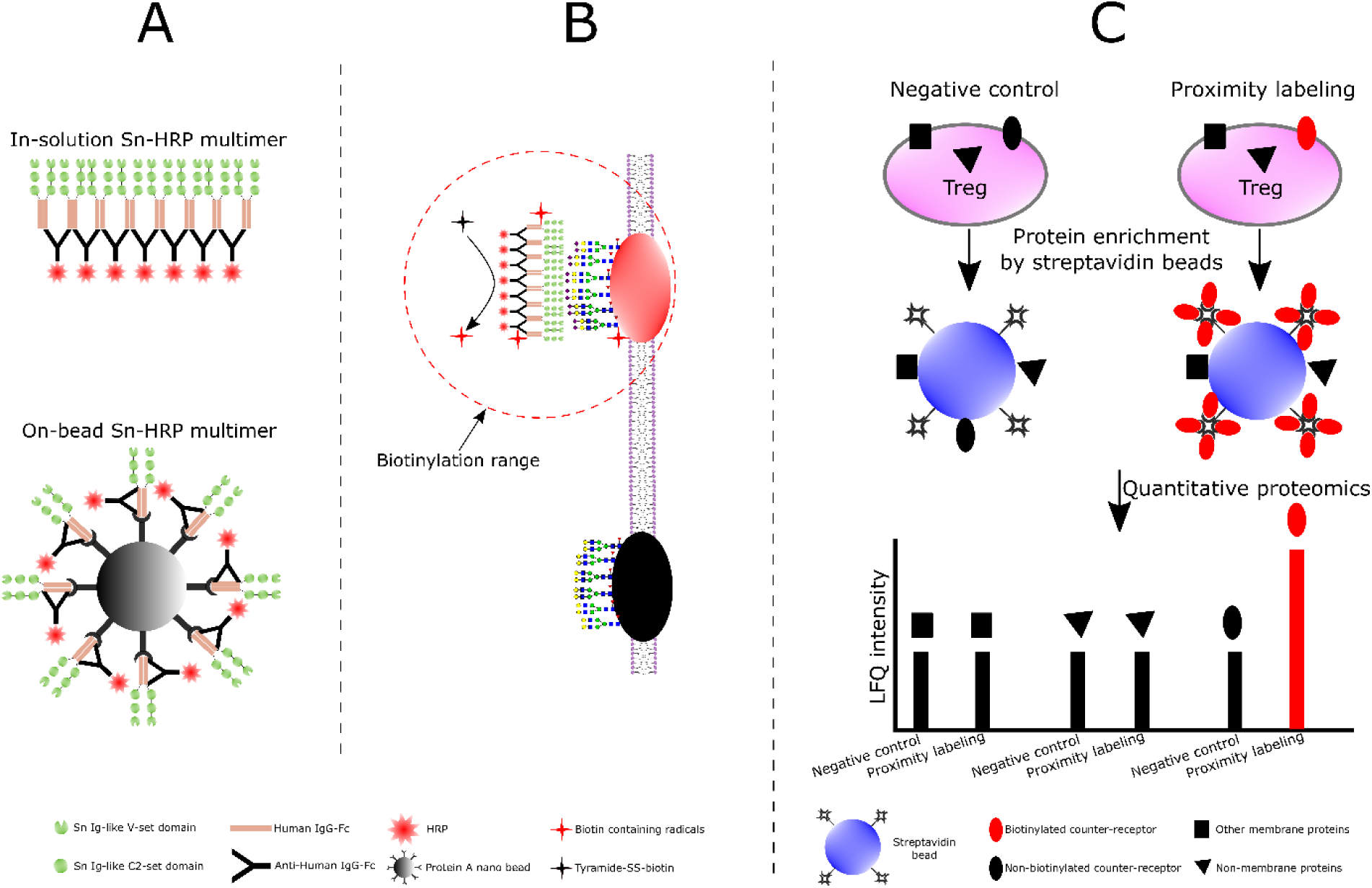
Design of the proximity labelling experiments. (**A)** Preparation of two types of Siglec-1-HRP (Sn-HRP) multimers. The in-solution multimers were prepared by mixing Siglec-1-huIgG Fc chimera (Sn-Fc) with HRP-conjugated polyclonal anti-huIgG Fc, while the on-bead multimers were prepared by immobilizing Siglec-1-huIgG Fc chimera (Sn-Fc) and HRP conjugated polyclonal anti-huIgG Fc on 50 nm protein A beads. **(B)** Mechanism of proximity labelling. After Siglec-1-HRP multimers bind to Siglec-1 counter-receptors on Treg membrane, tyramide-SS-biotin and H_2_O_2_ were added. In the presence of HRP, generation of short-range biotin radicals results in biotinylation of proteins in the immediate vicinity of the multimer. Siglec-1R97A-huIgG Fc (SnR97A-Fc) was used a negative control. **(C)** Identification of Siglec-1 counter-receptors. Only biotinylated proteins (coloured in red) can be selectively enriched by streptavidin beads and show significant quantitative changes in LFQ intensities in label free quantitative proteomic analysis.

Three biological replicates of activated Tregs were prepared for independent proximity labelling experiments using both in-solution and on-bead multimers, resulting in 6 data sets. For each data set, the log2 fold change of each protein relative to its negative control was calculated and visualized according to its cellular compartmentalisation (Figure 9A). A subset of membrane proteins with higher log2 fold changes were observed in the histogram, resulting in a small but clear tail, which we describe as a ‘proximity labeling tail’. This tail was only found for membrane proteins but not for cytoplasmic, nuclear or other proteins and suggests that only certain membrane proteins were biotinylated and selectively enriched. Five out of the 6 data sets showed the proximity labelling tail (Supplementary Figure 11) and these were selected for statistical analysis using volcano plots (Figure 9B). A small subset of proteins was found to have significant log2 fold changes. As expected, these were mainly membrane proteins.

**Figure 9.**
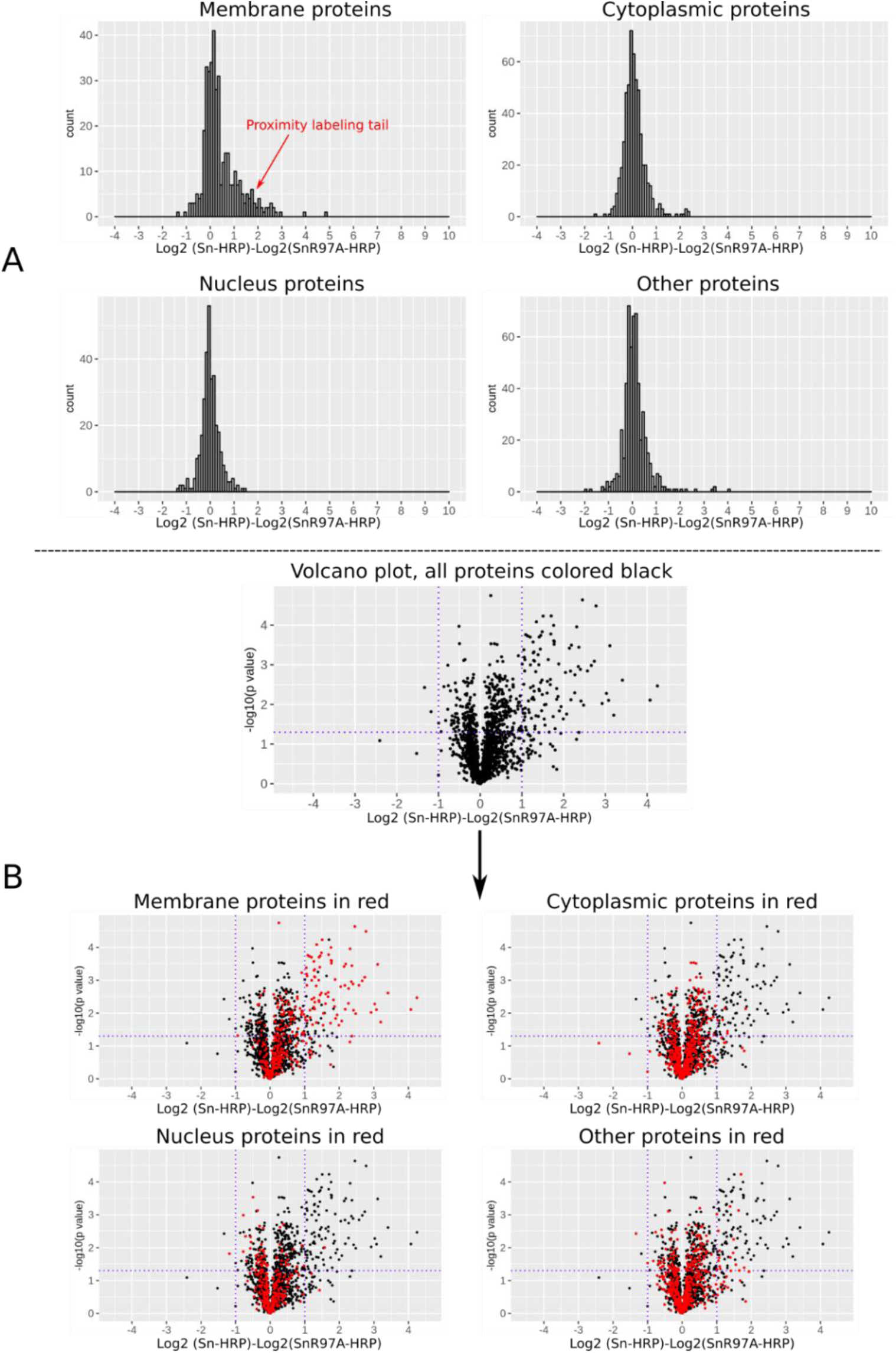
Quantitative proteomic analysis of proteins from proximity labelling experiments. After proximity labelling, the cells were lysed and streptavidin beads were used to enrich biotinylated proteins for proteomic analysis. **(A)** Histogram of log2 fold changes of proteins between a proximity labelling experiment and its negative control. Only membrane proteins contained a subset with higher log2 fold changes, resulting in a small proximity labelling tail. **(B)** Volcano plots of the proteomic data shown as log2 fold change and −log10 p values from paired t-test. A small subset of proteins was found to have significant log2 fold changes, which were mainly membrane proteins.

The glycosylated proteins from the significant hits on the volcano plot were selected for further data filtering; they were mapped back to the individual histogram of total membrane proteins (Supplementary Figure 12), and those which were outside the proximity labelling tail were filtered out. The final Siglec-1 counter-receptor list of 49 membrane proteins is shown in Table 1. We successfully identified proteins that make up the Siglec-1-HRP multimer complex, namely HRP, human IgG Fc and Siglec-1, with gene names PRXC1A, IGHG1 and Siglec1 respectively. We also identified CD43 which was proposed previously to function as a counter-receptor for Siglec-1 (van den Berg et al., 2001). These identifications serve as a proof-of-principle for the proximity labelling approach. Interestingly, Siglec-1 counter-receptors included a wide range of glycoproteins involved in a variety of functions, including the regulation of T cell activation and proliferation, such as CD80, CD200, CD69, CD150, PD-1 and PD-L1, adhesion molecules like CD166 and integrins, the IL-2 receptor (CD25) and transporters like the L-type amino acid transporter, 4F2 (Supplementary Table 2).

**Table 1.**
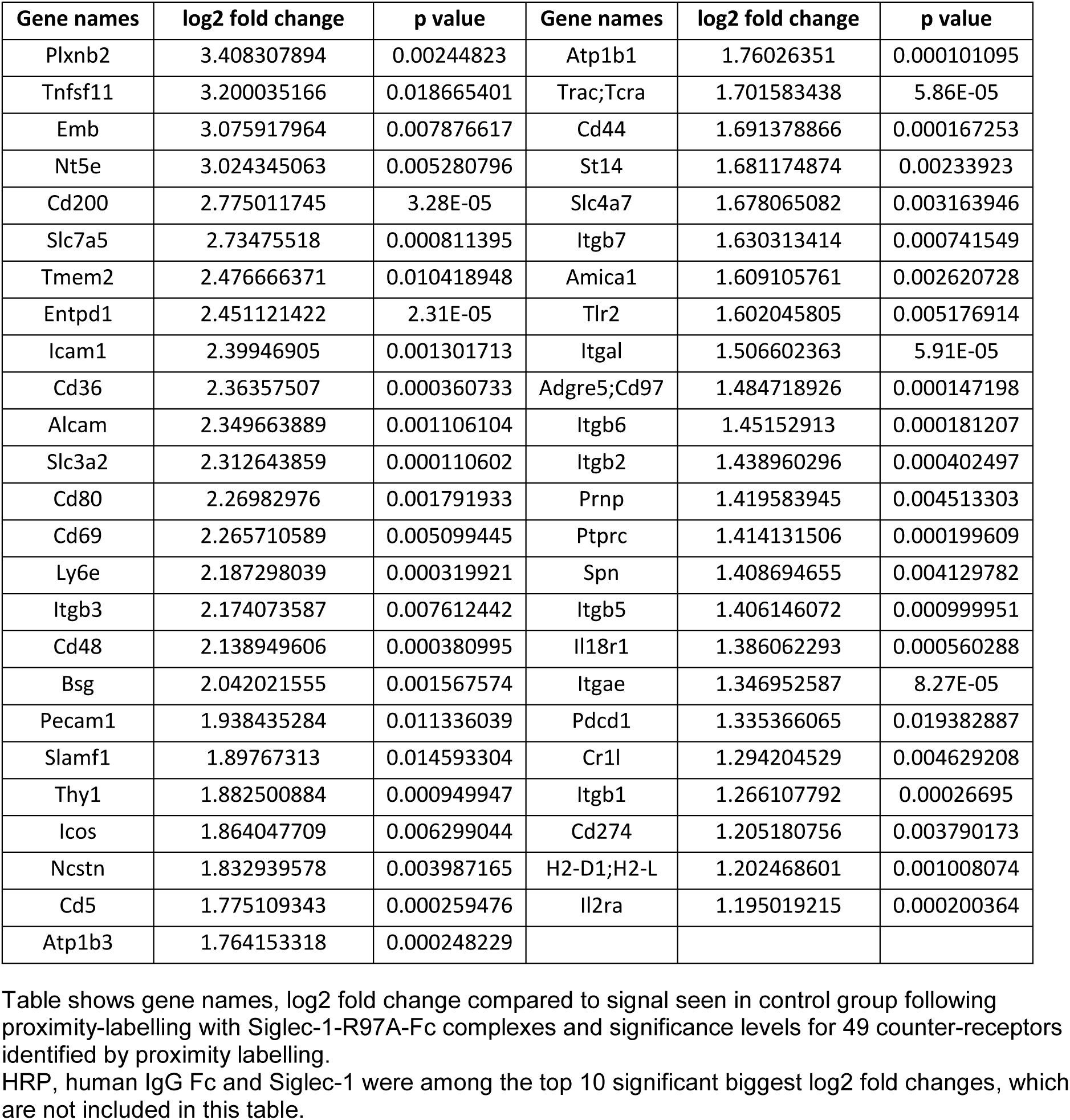
List of Siglec-1 counter-receptors identified by proximity labelling

### Characterization of Siglec-1 counter-receptors

To determine the proportion of membrane proteins constituted by the 49 Siglec-1 counter-receptors, proteomics analyses were undertaken on activated Tregs. To maximise the number of membrane proteins identified, two approaches were used, either by performing proteomics of whole cell lysates or following cell surface biotinylation and enrichment of labelled proteins with streptavidin beads. This combined approach led to identification of 943 membrane proteins, suggesting that the 49 Siglec-1 counter-receptors comprise approximately 5% of the total membrane proteins on activated Tregs (Figure 10A).

**Figure 10.**
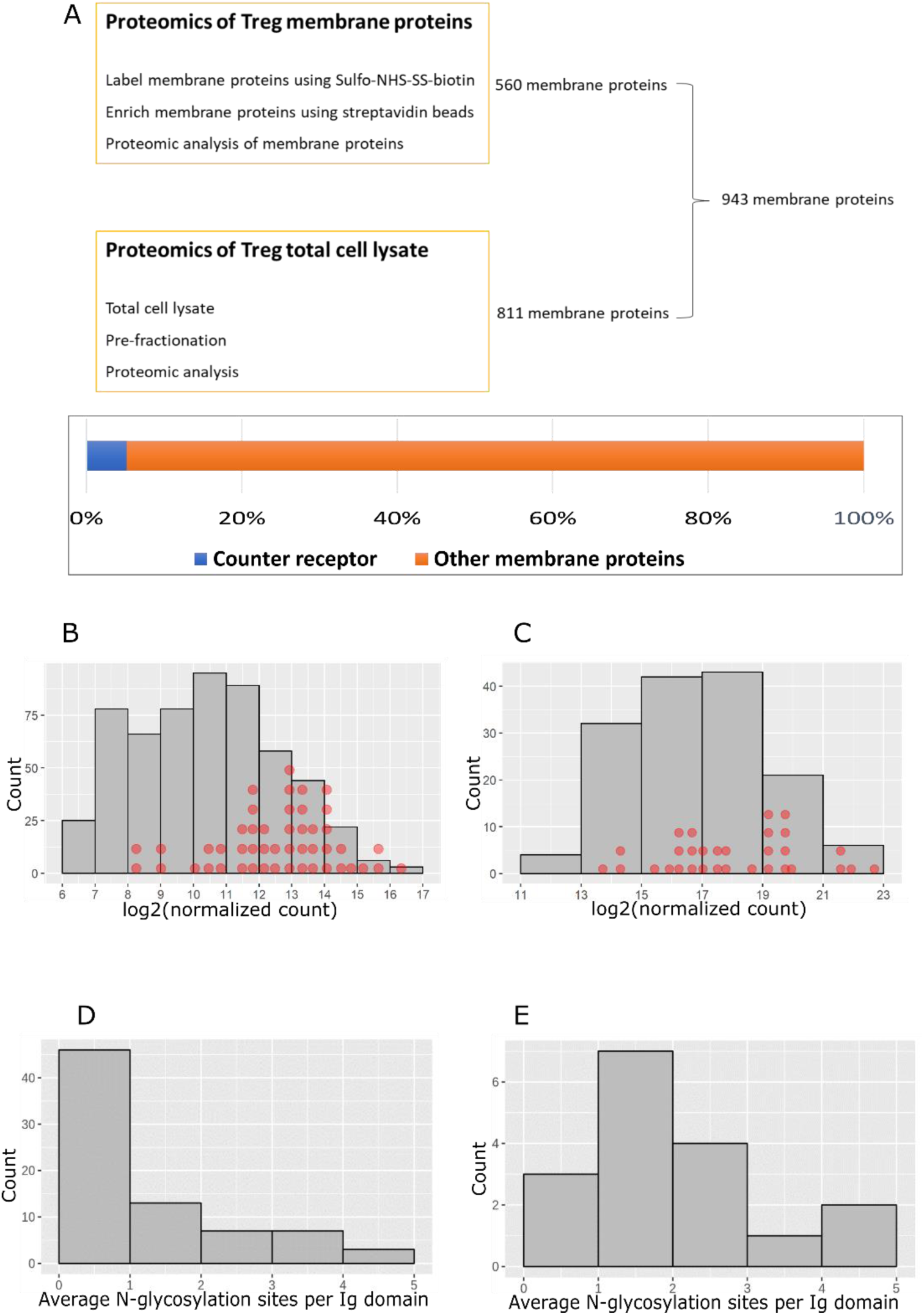
Characterization of Siglec-1 counter-receptors. **(A)** Total membrane proteins on activated Tregs were identified by combining the data from proteomics of total cell-surface proteins and proteomics of total cell lysates. Membrane proteins identified in at least 2 of 3 biological replicates were selected. Siglec-1 counter-receptors make up 5.2% of the total membrane proteins. **(B)** Counter-receptor mRNAs were mapped to the mRNAs of membrane glycoproteins on activated Tregs. The mRNAs with normalized counts above 100 were used for the histogram plot. Each red dot represents a counter-receptor. **(C)** Siglec-1 counter-receptors were mapped to the histogram of copy number per cell of membrane glycoproteins on activated Tregs. Each red dot represents a counter-receptor. **(D)** Predicted N-glycosylation sites per Ig like domain for membrane glycoproteins which are not Siglec-1 counter-receptors. **(E)** Predicted N-glycosylation sites per Ig like domain for Siglec-1 counter-receptors.

We next asked whether the Siglec-1 counter-receptors were distributed amongst the more abundant membrane glycoproteins using both the RNA-Seq and proteomics datasets (Figure 10B,C). This revealed that Siglec-1 counter-receptors were distributed across the range of proteins from low to high abundancy, but were more enriched amongst the high abundancy proteins. To investigate whether Siglec-1 counter-receptors might be more heavily glycosylated than other membrane proteins, we compared 16 counter-receptors belonging to the Ig superfamily with 56 non-counter-receptors of the Ig superfamily identified by proteomics. The results showed that most of the counter-receptors had more than 1 predicted N-glycosylation site per Ig-like domain, while most of the non-counter-receptors had less than 1 site per domain (Figure 10 D,E). This suggests that glycosylation density is an important determinant of functioning as a Siglec-1 counter-receptors, presumably by allowing them to mediate higher avidity binding to clustered Siglec-1. Next, we asked if the counter-receptors were upregulated on activated Tregs compared to resting cells. This was investigated at both the protein level using flow cytometry (Figure 11A) and at the mRNA level using RNA-Seq data (Figure 11B). Several proteins, including CD80, PD-1 and CD274 (PD-L1), showed increased protein expression in activated Tregs, which correlated with increased mRNA levels, but for about half of the counter-receptors, mRNA levels were either not changed or were decreased (Figure 11B).

**Figure 11.**
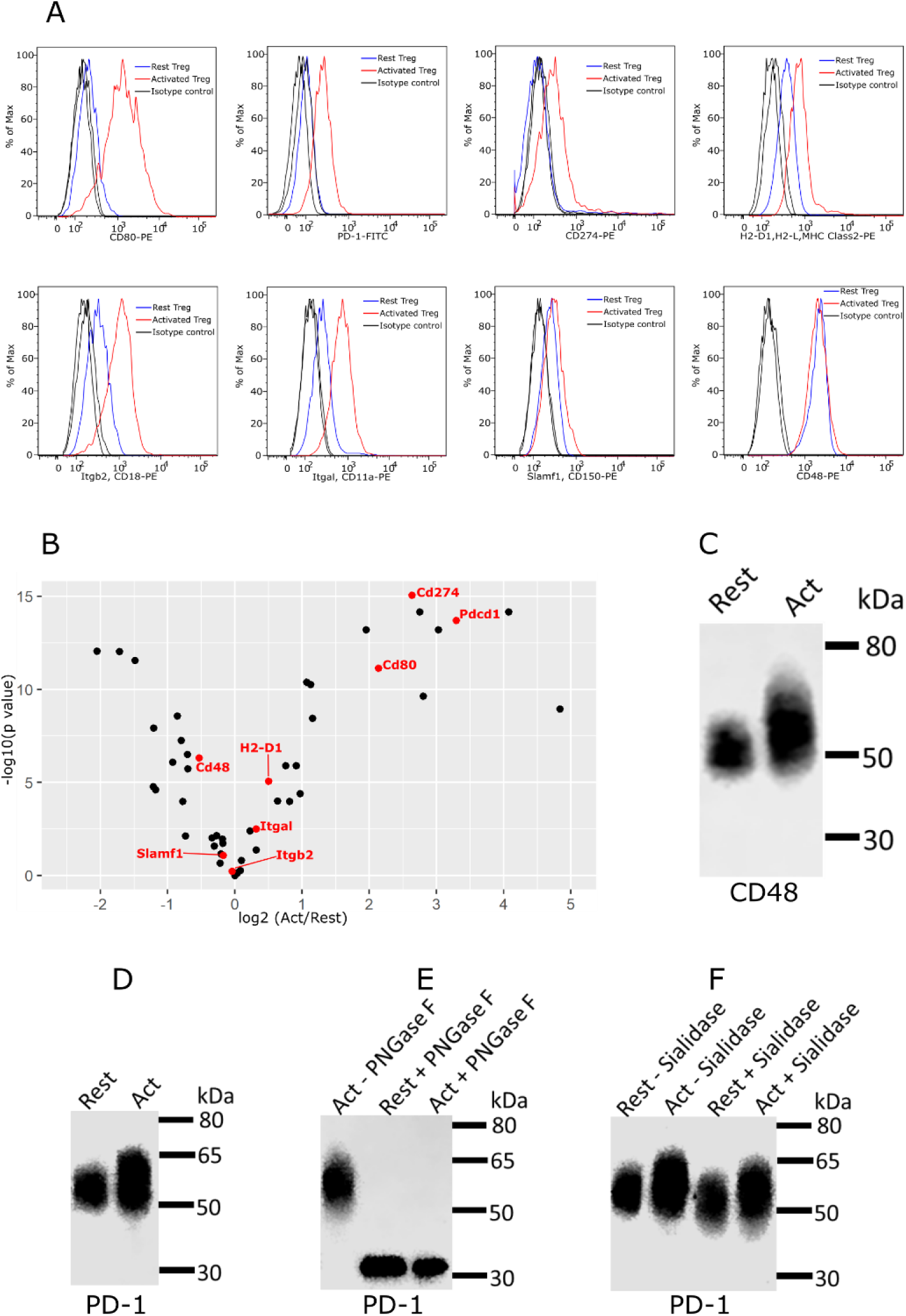
Expression and glycosylation of Siglec-1 counter-receptors on resting and activated Tregs. **(A)** Eight Siglec-1 counter receptors were randomly selected for cytometry analysis. All experiments were performed 3 times and similar results were observed. **(B)** Normalized mRNA counts of Siglec-1 counter receptors were obtained from Treg RNA-Seq data. The result is visualized in a volcano plot. Only a small subset of counter-receptors showed strongly increased gene expression following Treg activation. The counter-receptors selected for flow cytometry analysis are highlighted in red. **(C)** Western blot showing that CD48 had a higher molecular weight in activated Tregs. **(D)** Western blot of PD-1 affinity-purified from resting and activated Tregs.. PD-1 showed a dramatic decrease of molecular weight after PNGase F digestion **(D)**, PD-1 from activated Tregs had higher molecular weight **(E)** and following sialidase treatment, PD-1 from resting and activated Tregs migrated at a slightly reduced molecular weight indicating the presence of sialic acids **(F)**.

As a first step to investigate glycosylation changes in Siglec-1 counter-receptors, we performed western blotting on CD48, as an example of a protein that did not change expression, and PD-1, as an example of a protein that is upregulated on activated Tregs. Compared to resting Tregs, both CD48 and PD-1 from activated Tregs displayed a more heterogeneous smear and increased MW by SDS-PAGE, indicating increased glycosylation (Figure 11C,D). For PD-1, this was confirmed following treatment of affinity-purified PD-1 with PNGase F to remove N-glycans which resulted in PD-1 from resting and activated Tregs migrating similarly by SDS-PAGE (Figure 11E). Finally, affinity-purified PD-1 from resting and activated Tregs was treated with sialidase, leading to increased migration by SDS-PAGE showing that it carries sialylated glycans with potential to be recognised by Siglec-1 (Figure 11F).

## Discussion

The major aim of this study was to obtain insights into the molecular basis of Siglec-1-dependent interactions of macrophages with Tregs and thus improve our understanding of how this lectin promotes inflammatory responses in certain autoimmune diseases. Consistent with *in vivo* and *in vitro* studies (Kidder et al., 2013; C. Wu et al., 2009), we found Siglec-1 binding to Tregs depended completely on Treg activation. Based on this finding, a key part of our strategy was to analyse glycosylation changes by performing side-by-side comparisons of resting and activated Tregs using a range of unbiased and targeted approaches. These included RNA-Seq, glycomics, proteomics and staining with lectins and anti-glycan antibodies. Furthermore, we used a recently-described proximity labelling strategy (Chang et al., 2017; G. Wu et al., 2017) to identify membrane proteins on activated Tregs that could function as Siglec-1 counter-receptors. This comprehensive combined approach has provided a wealth of new information regarding glycomics changes and Siglec-1 recognition following Treg activation. Clear changes in glycosylation were observed for both N-glycans and glycolipids and many glycogenes were altered in expression following Treg activation. Although the precise mechanisms by which these changes lead to increased Siglec-1 binding remain to be determined, they are likely to involve complex protein- and lipid-dependent modification of sialylation at an individual carrier level. The identification of 49 Siglec-1 glycoprotein counter-receptors provides important clues as to how Siglec-1 can mediate adhesion and signalling to activated Tregs. These counter-receptors were dominated by adhesion and signalling glycoproteins mainly related to T cell activation and proliferation (Table 1, Supplementary Table 2). Importantly, several counter-receptors showed increased expression at the protein level and preliminary analyses showed increased glycosylation following Treg activation. The combined effect of all changes noted above could lead to increased Siglec-1 binding and signalling to activated Tregs.

Although the focus of this study was on the lectin Siglec-1, there are many other endogenous lectins such as galectins and C-type lectins whose binding could be greatly affected by the glycosylation changes observed leading to altered Treg functions (van Kooyk & Rabinovich, 2008). The glycan remodelling of Tregs could also be important in intrinsic functions of these cells since it is well known that glycosylation has complex pleiotropic effects on cell function and behaviour (Varki, 2017).

Previous studies with effector CD4 T cells have also demonstrated activation-dependent increases in Siglec-1 binding, and these were ascribed to a switch from NeuGc to NeuAc and increased α2,3-linked sialic acids following activation (Naito-Matsui et al., 2014; Redelinghuys et al., 2011). In the present study, we were unable to see obvious global changes in either of these parameters using anti-NeuGc antibody staining and MALII plant lectin that binds α2,3 linked sialic acids. The unaltered expression of NeuGc at the cell surface was consistent with RNA-Seq analysis of *Cmah*, which encodes the enzyme responsible for converting CMP-NeuAc to CMP-NeuGc, did not show a significant change of expression in resting and activated Tregs. It appears, therefore, that regulation of Siglec-1 ligand expression by Tregs is fundamentally different from effector CD4 T cells. However, glycomics analysis did identify minor changes in NeuGc capped glycans upon Treg activation. We also detected clear reductions in NeuGc expression in glycolipid analysis following activation, but as these gangliosides are likely to be relatively minor carriers of sialic acids compared to glycoproteins, their altered levels are probably not detectable with anti-NeuGc antibody and their impact on Siglec-1-dependent binding is unclear. Interestingly, a previous study has found that ST6Gal I showed 4-7 times higher affinity for CMP-NeuGc than CMP-NeuAc, while ST3Gal I showed no significant difference between them (Hamamoto, Kurosawa, Lee, & Tsuji, 1995). This implies that sialyltransferases can have different activities towards CMP-NeuGc and CMP-NeuAc, which could lead to different incorporation of NeuGc and NeuAc to target glycans. It is possible that the specific changes of NeuGc capped glycans were due to altered sialyltransferase gene expression upon Treg activation.

While there was no overall change in levels of α2,3-sialylation measured with MALII lectin in activated Tregs, RNA-Seq showed reduced expression of sialyltransferases *St3gal1, 2* and *6* and increased expression of *St3gal5*. These observations suggest that individual glycoconjugates that function as acceptors for these enzymes may differ significantly in α2,3 sialylation between resting and activated Tregs. For example, expression of *St3gal6* has been associated with generation of the selectin ligand sialylLe^x^ (Yang, Nussbaum, Grewal, Marth, & Sperandio, 2012) and its reduced expression in activated Tregs could be important for modulating homing of these cells to inflamed sites. ST3Gal5 is also known as GM3 synthase which converts lactosyl ceramide to GM3. GM3 can be further converted to α2,8-disialylated GD3 by ST8Sia1 (GD3 synthase). The increased expression of *St3gal5* and decreased expression of *St8sia1* suggests that GM3 is upregulated on activated Tregs where it could function as a ligand for Siglec-1. GM3 is known to be a dominant ligand for Siglec-1 interactions with retroviruses such as HIV and is able to mediate recognition and internalisation of viral particles by Siglec-1-expressing macrophages (Izquierdo-Useros et al., 2014; Puryear et al., 2013). While future studies are required to investigate expression of GM3 following Treg activation and its potential role in Siglec-1 recognition, we were unable to detect GM3 in the glycolipid analysis. This may be due to the low level of *St3Gal5* expression relative to other *St3Gals*, resulting in synthesis of GM3 at levels below our detection threshold.

An important aim of this study was to identify glycoproteins on activated Tregs that could function as counter-receptors for Siglec-1 and mediate the biological effects, namely suppression of Treg expansion during autoimmune inflammatory responses. Although pulldown approaches are widely used to identify high affinity protein binding partners, these may not be reliable for identifying counter-receptors for lectins such as Siglec-1 which rely on high avidity interactions with clustered glycan ligands. Clustering could result from organisation of membrane glycoproteins and glycolipids within microdomains of the membrane, as well as through a high density of glycan ligands on individual glycoproteins. Pull-down approaches depend on detergents to lyse cells and solubilise membrane proteins. As such, they disrupt the organization and clustering of glycoproteins in the cell membrane that may be critical for Siglec-1 binding.

Proximity labelling is a recently developed method, which has been used widely to study protein-protein interactions in living, intact cells (Branon et al., 2018; Lam et al., 2015; Li et al., 2014; Rees, Li, Perrett, Lilley, & Jackson, 2015, 2017; Roux, Kim, Raida, & Burke, 2012). In general, this approach uses enzyme conjugates as baits that bind cellular target proteins. Addition of suitable substrates leads to the generation of highly reactive radicals that tag neighbouring proteins which can then be enriched and identified by mass spectrometry. This method can also be used to identify Siglec counter-receptors by building Siglec-HRP multimers. We previously prepared in-solution Siglec-1-HRP multimers by mixing Siglec-1-huIgG-Fc fusion protein with HRP-conjugated polyclonal anti-huIgG-Fc and successfully identified glycophorin as a Siglec-1 counter-receptor on human erythrocytes (G. Wu et al., 2017). Another type of in-solution Siglec-HRP multimer was independently developed by another group which identified CD22 counter-receptors on B cells and Siglec-15 counter-receptors on RAW264.7-derived osteoclasts (Chang et al., 2017). In this study, we used two types of multimer for proximity labeling: in-solution multimers and on-bead multimers (Figure 8). Glycoproteins identified by both types of multimer were selected as Siglec-1 counter-receptors.

An initial characterization of Siglec-1 counter-receptors revealed they represent 5% of the total membrane proteins and tend to be amongst the more abundantly expressed membrane glycoproteins. We did not see evidence for strong enrichment of any minor proteins which might be expected if Siglec-1 had high affinity binding partners requiring protein-protein interactions as well as protein-glycan interactions. Although most lectins rely solely on high avidity glycan interactions for protein binding, P-selectin is an example of a lectin that can use additional, non-glycan interactions to mediate high affinity binding to P-selectin glycoprotein ligand-1 (McEver & Cummings, 1997). Consistent with a glycan-dependent, protein-independent mode of interaction to clustered glycans, Siglec-1 counter-receptors carry a higher density of predicted N-glycosylation sites than non-counter-receptors. O-glycosylation could also be important for Siglec-1 binding, especially for mucin-like proteins that contain a high density of O-glycans. Previously, using a pull-down approach, the mucin-like proteins, CD43 and Muc-1, were identified as Siglec-1 counter-receptors on a T cell line and breast cancer cells respectively (Nath et al., 1999; van den Berg et al., 2001). In the present study, we also identified CD43 as a Siglec-1 counter-receptor on activated Tregs, alongside 48 other glycoproteins. Unlike N-glycosylation, O-glycosylation does not have consensus sequences, making it more difficult to compare O-glycan densities on the counter-receptors versus non-counter-receptors.

Identification of potential binding partners allowed us to ask if these were upregulated on activated Tregs which could then help explain the increased binding of these cells to Siglec-1 compared to resting Tregs. Indeed, flow cytometry of several counter-receptors showed increased expression following Treg activation but there were clear exceptions such as CD48 whose expression remained unchanged. RNA-Seq data also indicated that the expression of many Siglec-1 counter-receptors was either unaltered or even reduced following Treg activation. Even if expression of counter-receptors is not changed, increased glycosylation and sialylation following activation could lead to enhanced Siglec-1 binding. In support of this possibility, KEGG pathway analysis of the RNA-Seq data indicated that Treg activation was accompanied by increased flux through the N-glycosylation pathway. In particular, activated Tregs had enhanced expression of several components of the OST complex which can be rate-limiting for N-glycosylation of proteins (Rinis et al., 2018). Direct evidence for increased glycosylation of counter-receptors following Treg activation was seen for CD48 and PD-1, both of which showed an increase in apparent MW and in the case of PD-1 this was shown to be sensitive to PNGase F that specifically cleaves N-glycans. Further work is required to investigate the glycosylation changes in more detail and determine if these lead to increased Siglec-1 binding for individual counter-receptors. Glycoprotemics is a powerful strategy which can map glycan structures to each glycosylation site of a glycoprotein (Tissot et al., 2009). However, glycoproteomics requires relatively large amounts of purified target glycoproteins, which, for primary cells such as Tregs, is a major technical challenge (Plomp et al., 2014; G. Wu et al., 2016).

The glycan-dependent interaction of Siglec-1 on macrophages with the set of counter-receptors expressed on Tregs could trigger a range of biological activities, with the net outcome being suppression of Treg expansion, as observed previously (C. Wu et al., 2009). The identification of CD25/IL-2 receptor could be relevant in this regard since Tregs do not produce IL-2 but they require IL-2 for survival and proliferation (Horwitz, Zheng, Wang, & Gray, 2008). Modulation of CD25 function via Siglec-1 binding could therefore affect Treg cell expansion under inflammatory conditions. Of particular interest from this perspective are regulatory receptors such as PD-1 which is an important immune-inhibitory receptor. It can be exploited by pathogens and cancer cells to escape T-cell–mediated immune responses (Boussiotis, 2016). Antibodies targeting PD-1 and its ligands are used to treat human cancers in checkpoint immunotherapy (Chen & Han, 2015). PD-1 also plays a vital role in the maintenance of peripheral tolerance by thwarting autoreactive T cells (Francisco, Sage, & Sharpe, 2010). Two PD-1 ligands have been identified: PD-L1 (CD274) and PD-L2 (CD273). The results of our proximity labeling studies suggests Siglec-1 could act as an additional PD-1 ligand. PD-1 is a heavily glycosylated protein; both mouse and human PD-1 contain a single Ig-like domain with 4 N-glycosylation sites (Tan et al., 2017). As shown in the present study, deglycosylated PD-1 has a molecular weight of about 30 kDa, which can increase up to 65 kDa when glycosylated. Interestingly, the involvement of Siglec-1 in PD-1 signalling has been reported previously (Kirchberger et al., 2005). Human monocyte-derived dendritic cells treated with human rhinoviruses showed upregulated expression of PD-L1 and induced expression of Siglec-1. These dendritic cells had an inhibitory phenotype, which diminished their capacity to stimulate alloreactive T cells and induced a promiscuous and deep T cell anergy. The inhibitory phenotype of these dendritic cells could be reversed by blocking both PD-L1 and Siglec-1 (Kirchberger et al., 2005). In contrast to the function of PD-1 in effector T cells, PD-1 is important for Treg development and activity (Gianchecchi & Fierabracci, 2018). The impact of Siglec-1 on PD-1 function in T cells is an important area for future studies.

In conclusion, we provide the first comprehensive analysis of glycan changes in activated Tregs that lead to recognition by the macrophage lectin, Siglec-1. We furthermore provide insights into glycoprotein counter-receptors expressed by these cells that are likely to be important in mediating the biological functions of Siglec-1, by promoting inflammatory responses via suppression of Treg expansion during autoimmune disease of the nervous system.

## Materials and Methods

### Generation and culture of Tregs

RPMI 1640 medium with L-glutamine (Gibco™), FBS (Gibco™), Penicillin-Streptomycin (Gibco™), 2-mercaptoethanol (Gibco™), functional grade anti-mouse CD28 (eBioscience™, clone: 37.51), functional grade anti-mouse IL-4 (eBioscience™, clone: 11B11), functional grade anti-mouse INF-γ (eBioscience™, clone: XMG1.2) were from ThermoFisher, Loughborough, UK. Anti-mouse CD3 (clone: 145-2C11) was from Biolegend, London, UK. CD4+ T Cell Isolation Kit was from Miltenyi Biotec Ltd., Surrey, UK. Mouse IL-2 and human TGF-β were from PeproTech, London, UK. TPP® 12 well plate was from Merck, Dorset, UK.

Mouse T cells were isolated from spleen and lymph nodes using CD4+ T Cell Isolation Kits following the supplier’s protocol. Mouse iTregs were generated and grown in 12-well plates. The wells were coated with 1 ml PBS per well containing 10 ug/ml anti-CD3 for 2 hours at 37 °C. The non-bound antibody was removed by washing the plate twice using PBS. The isolated cells were suspended at a concentration of 1-2 × 10^6^/ml in culture medium which is RPMI 1640 with L-glutamine, 10% FBS, 100 U/ml Penicillin-Streptomycin, and 50 uM 2-mercaptoethanol. The cells were induced for 4-5 days at 1 ml of cell suspension per well with 2 ug/ml functional grade anti-CD28, 10 μg/ml functional grade anti-IL-4, 10 μg/ml functional grade anti-INF-γ, 20 ng/ml IL-2 and 5 ng/ml TGF-β. Fresh culture medium, anti-CD28, anti-IL-4, anti-INF-γ, IL-2, and TGF-β were supplemented when the cell culture medium became yellow. After induction, the cells were washed twice with culture medium. The cells were re-suspended at a concentration of 10^6^/ml in the culture medium and expanded for 4 days with 20 ng/ml IL-2 and 5 ng/ml TGF-β. Fresh culture medium, IL-2 and TGF- β were supplemented when the culture medium became yellow. To obtain activated Tregs, the expanded Tregs were suspended at a concentration of 10^6^/ml in culture medium with 20 ng/ml IL-2 and 5 ng/ml TGF-β, and were cultured at 1 ml cell suspension per well in 12 well plate pre-coated with different concentrations of anti-CD3 in PBS. The anti-CD3 concentration for activated Tregs for proximity labelling, proteomics and RNA-Seq was 0.125 μg/ml. Tregs were activated for 3 days for proximity labelling and protemics experiments, and for 36 hours for RNA-Seq experiments.

### Flow cytometry

Anti-mouse CD4-PE-Cy7 (eBioscience™, clone: GK1.5), anti-mouse Foxp3-APC (eBioscience™, clone: FJK-16s), anti-mouse PD-1-FITC (clone J43), anti-mouse MHC Class II-PE (eBioscience™, clone: AF6-120.1), anti-mouse CD150-PE (eBioscience™, clone: mShad150),Fixable Viability Dye eFluor™ 450 (eBioscience™), Foxp3 / Transcription Factor Staining Buffer Set, HyClone™ FetalClone™ II serum were from ThermoFisher, Loughborough, UK. Anti-mouse CD80-PE (clone: 16-10A1), anti-mouse CD274-PE (clone: MIH7), anti-mouse CD18-PE (clone: M18/2), anti-mouse CD11a-PE (clone: I21/7), anti-mouse CD48-PE (HM48-1), streptavidin-FITC, anti-Neu5Gc antibody kit were from Biolegend, London, UK. FITC conjugated goat anti-human IgG Fc was from Merck, Dorset, UK. Biotinylated plant lectin MAL II and SNA were from Vector® Laboratories, Peterborough, UK. Mouse Siglec-1-human IgG Fc chimera was expressed using stably-transfected Chinese Hamster Ovary cells (Kidder et al., 2013).

Before staining for the presence of antigens and glycans, cells were labelled with the Fixable Viability Dye eFluor™ 450, following the supplier’s instructions, and live cells were gated for flow cytometry analysis. All antigen and glycan staining steps were carried out on ice. For staining Foxp3 and NeuGc, Foxp3 / Transcription Factor Staining Buffer Set and the buffer from anti-Neu5Gc antibody Kit were used as staining buffer and washing buffer respectively. For the other staining experiments, 1% HyClone™ FetalClone™ II serum in PBS was used as staining buffer, antibody diluting buffer, washing buffer and cell storage buffer. Foxp3 staining and NeuGc staining were done according to the kit supplier’s instructions. For staining antigens, the cells were washed with staining buffer, stained on ice for 30 min, and non-bound washed for flow cytometry analysis. For staining Siglec-1 ligands, Siglec-1-human IgG Fc chimera was mixed with FITC-conjugated goat anti-human IgG Fc at a final concentration of 1 μg/ml in staining buffer, and incubated on ice for 30 min to prepare the pre-complex. The pre-complex was used according to the antibody staining procedure to stain glycan ligands for flow cytometry analysis. Biotinylated MAL II and SNA staining also followed the antibody staining procedure. and detected by flow cytometry using streptavidin-FITC. All flow cytometry data were analysed using FlowJo. A Siglec-1-Fc mutant (SnR97A-Fc) was used as a negative control for Siglec binding experiments. The chicken IgY isotype in the anti-Neu5Gc antibody Kit was used as a negative control for NeuGc staining experiments. Streptavidin-FITC was used as a negative control for MAL II and SNA binding experiments.

### Proximity labelling

Protein A beads and µ columns were from Miltenyi Biotec Ltd., Surrey, UK. Tyramide-SS-biotin was from Iris Biotech GmbH, Marktredwitz, Germany. HRP conjugated goat anti-human IgG Fc was from Abcam, Cambridge, UK. Catalase was from Merck,Dorset, UK. H_2_O_2_ was from VWR, Leicestershire, UK.

Proximity labelling was done using activated Tregs induced from independent experiments, and the gap between these experiments was set no less than 2 weeks. All proximity labelling steps were carried out on ice. In-solution Siglec-1-HRP multimers were prepared by incubating Siglec-1-human IgG Fc chimera with HRP-conjugated goat anti-human IgG Fc at a final concentration of 1 μg/ml and 3 μg/ml in labelling buffer (1%HyClone™ FetalClone™ II serum in PBS) for 30 min. 5×10^6^ Tregs were washed twice with labelling buffer, mixed with 300 μl Siglec-1-HRP multimer and incubated for 30 min.

For proximity labelling using on-bead Siglec-1-HRP multimer, 270 μl 10 μg/ml Siglec-1-human IgG Fc chimera was incubated with 30 μl Protein A nanobeads for 60 min. The non-bound were removed by washing the beads on a µ column. The beads were eluted from the µ column using 300 μl labelling buffer, mixed with 5×10^6^ Tregs and incubated for 30 min. HRP-conjugated goat anti-human IgG Fc was added to the cells to a final concentration of 3 μg/ml and incubated for another 30 min.

HRP substrate, Tyramide-SS-biotin and H_2_O_2_, were added to a final concentration of 95 µM and 0.01% respectively. The sample was left on ice for 2 min and the reaction was quenched by washing 3 times with 3 ml 100 U/ml catalase in labelling buffer.

### Glycomics

PNGase F (cloned from Flavobacterium *meningosepticum* and expressed by *E.coli*), CHAPS and DTT were from Roche, Welwyn Garden City, UK. Sep Pak C18 Cartridges were from Waters, London, UK. Slide A Lyzer ® dialysis cassettes (3.5 kDa molecular weight cut off), TMT10plex™ Isobaric Label Reagent Set, TCEP, TEAB were from ThermoFisher, Loughborough, UK. Protein DeglycosylationMix II Kit was from New England Biolabs, Hitchin, UK. LysC was from Alpha Laboratories Ltd, Eastleigh, UK. Trypsin was from Promega, Southampton. UK. Paramagnetic bead (SP3 bead) and other general chemical reagents were from Merck, Dorset, UK.

Glycomic analysis of glycoproteins was done following the protocol published previously (Jang-Lee et al., 2006). Briefly, Tregs were homogenized by sonication in 25 mM Tris, 150 mM NaCl, 5 mM EDTA, and 1% CHAPS, pH 7.4, dialysed in dialysis cassettes, reduced by DTT, carboxymethylated by IAA, and digested by trypsin. N-glycans were removed from glycopeptides by PNGase F, which were then isolated by Sep Pak C18 Cartridges and permethylated for mass spectrometry analysis. Glycomic analysis of glycolipids was done following a previous protocol (Parry et al., 2007). Glycomic data were analysed using Data Explorer™ version 4.6 from AB Sciex.

### Proteomics

Proteomic sample preparation was done using paramagnetic bead (SP3 bead) technology following the protocol published previously (Hughes et al., 2014). For histone ruler proteomics, the cells were lysed and sonicated in cell lysis buffer (4% SDS, 10 mM TCEP, 50 mM TEAB in H2O). The proteins were then alkylated using iodoacetamide, cleaned on SP3 beads, and digested to peptides using trypsin and LysC. The peptides were TMT labelled according to the supplier’s instructions, cleaned on SP3 beads, eluted and fractionated by high pH reversed phase chromatography for mass spectrometry analysis. For Siglec-1 counter-receptor identification and membrane proteomics, the biotinylated proteins were enriched and cleaned on streptavidin beads. The glycans were removed using Protein Deglycosylation Mix II under Non-Denaturing Reaction Conditions following the protocol from the supplier. The samples were cleaned on streptavidin beads, eluted using cell lysis buffer and processed using paramagnetic bead (SP3 bead) technology for label free proteomic analysis.

The proteomic raw data were imported to MaxQuant Version 1.6.2.3 (Tyanova, Temu, & Cox, 2016) to search protein FASTA files of mouse, human IgG Fc and HRP in Uniprot database. The LFQ intensities of proteins were used for downstream analysis (Cox et al., 2014) by Perseus Version 1.6.0.7 (Tyanova, Temu, Sinitcyn, et al., 2016) and R Version 3.6.3. The R scripts were uploaded to GitHub (https://github.com/fromgangwu/R-scrip-for-Treg-data-analysis.git). Gene Ontology Cellular Component analysis and glycosylation analysis were based on the reviewed mouse protein entries in Uniprot database. Siglec-1 counter receptors were identified through 4 consecutive steps of data filtration. The log2 fold change of the proteins in each sample was first visualized by histogram, and the samples without a proximity labelling tail were excluded. After that, the rest of the data were visualized in a volcano plot and the data points with significant changes were selected. Then, proteins without glycosylation or not located on plasma membrane were removed. Finally, the proteins were mapped back to each of the histogram of log2 fold change and proteins found outside the proximity labelling tail in any histogram were removed. For the identification of total membrane proteins, both total cell lysate and membrane enriched proteins from 3 biological replicates of activated Tregs were used for mass spectrometry analysis. The proteins that identified in at least 2 of 3 biological replicates were selected for Gene Ontology Cellular Component analysis to identify cell membrane proteins.

### RNA Sequencing and data analysis

Total RNA from rested and activated regulatory T cells (four biological replicates) were extracted using Qiagen RNeasy Mini kit according to the manufacturer’s instructions and quantified using the Qubit 2.0 Fluorometer (Thermo Fisher Scientific Inc, #Q32866) and the Qubit RNA BR assay kit (#Q10210). RNA sequencing libraries were prepared from 500 ng of each total-RNA sample using the NEBNEXT Ultra II Directional RNA Library Prep kit with Poly-A mRNA magnetic isolation (NEB #E7490) according to the manufacturer’s instructions.

Poly-A containing mRNA molecules were purified using poly-T oligo attached magnetic beads. Following purification the mRNA was fragmented using divalent cations under elevated temperature and primed with random hexamers. Primed RNA fragments were reverse transcribed into first strand cDNA using reverse transcriptase and random primers. RNA templates were removed and a replacement strand synthesised incorporating dUTP in place of dTTP to generate ds cDNA. The incorporation of dUTP in second strand synthesis quenches the second strand during amplification as the polymerase used in the assay is not incorporated past this nucleotide. AMPure XP beads (Beckman Coulter, #A63881) were then used to separate the ds cDNA from the second strand reaction mix, providing blunt-ended cDNA. A single ‘A’ nucleotide was added to the 3’ ends of the blunt fragments to prevent them from ligating to another during the subsequent adapter ligation reaction, and a corresponding single ‘T’ nucleotide on the 3’ end of the adapter provided a complementary overhang for ligating the adapter to the fragment. Multiple indexing adapters were then ligated to the ends of the ds cDNA to prepare them for hybridisation onto a flow cell, before 11 cycles of PCR were used to selectively enrich those DNA fragments that had adapter molecules on both ends and amplify the amount of DNA in the library suitable for sequencing. After amplification libraries were purified using AMPure XP beads.

Libraries were quantified using the Qubit dsDNA HS assay and assessed for quality and fragment size using the Agent Bioanalyser with the DNA HS kit (#5067-4626). RNA-Sequencing was carried out by The Genetics Core of Edinburgh Clinical Research Facility, University of Edinburgh using the NextSeq 500/550 High-Output v2 (150 cycle) Kit (# FC-404-2002) with a High Out v2.5 Flow Cell on the NextSeq 550 platform (Illumina Inc, #SY-415-1002). 8 libraries were combined in an equimolar pool based on the library quantification results and run across one High-Output Flow Cell. Sequencing resulted in paired-end reads 2 x 75 bp with a median of 50 million reads per sample.

The data were analysed by the Data Analysis Group, Division of Computational Biology, University of Dundee. The sequencing data were processed using a snakemake script in a conda environment. Reads were quality controlled using *FastQC* and *MultiQC*, mapped to GRCm38 assembly (Ensembl) of the mouse genome using *STAR* 2.6.1a (Dobin et al., 2013)and number of reads per gene was quantified in the same *STAR* run. Differentially expressed genes were quantified with *edgeR* v3.28.0 (Robinson, McCarthy, & Smyth, 2010). A Benjamini-Hochberg multiple test correction was applied to test P-values. The data reproducibility was tested using the distance matrix and clustering based on the read count per gene. The code used to process data is available at https://github.com/bartongroup/MG_GlycoTreg.

### Glycoprotein isolation, glycosidase digestion, Western blotting

Cyanogen bromide-activated Agarose, neuraminidase (*Vibrio cholerae*), and HRP-conjugated rabbit anti-goat antibody were from Merck, Dorset, UK. Monoclonal anti-mouse PD-1 (clone: J43) and NuPAGE SDS-PAGE system was from ThermoFisher,Loughborough, UK. Polyclonal goat anti-mouse PD-1, Polyclonal goat anti-mouse CD48 was from R&D systems, Abingdon, UK. PNGase F was from New England Biolabs, Herts UK.

For PD-1 analysis, anti-mouse PD-1 (clone: J43) was conjugated to cyanogen bromide-activated Agarose according to the supplier’s instructions. PD-1 from resting and activated Tregs were affinity purified using the anti-PD1 beads for glycosidase digestion and Western blot analysis. PNGase F digestion was done according to the supplier’s instructions. Sialidase digestion was done in 50 mM sodium acetate, 4 mM calcium chloride, pH 5.5, at 37 °C for 2 h. SDS-PAGE of PD-1 was done using the NuPAGE system, and the protein was transferred to PVDF membrane, blotted with goat anti-mouse PD-1 and HRP-conjugated rabbit anti-goat antibody. CD48 was blotted with goat anti-mouse CD48 and HRP-conjugated rabbit anti-goat antibody.

## Acknowledgements

We would like to thank the staff in our animal facility for their help in routine breeding and maintenance of mice. We thank the University of Dundee Proteomics Facility and the Flow Cytometry and Cell Sorting Facility at the University of Dundee for their assistance. This work was supported by Wellcome Trust Investigator Award (103744) to P.R.C. Glycomics analysis was supported by the Biotechnology and Biological Sciences Research Council, grant number BB/K016164/1.

