## Supplementary data for "Regulatory T cell activation triggers specific changes in glycosylation associated with Siglec-1-dependent inflammatory responses"

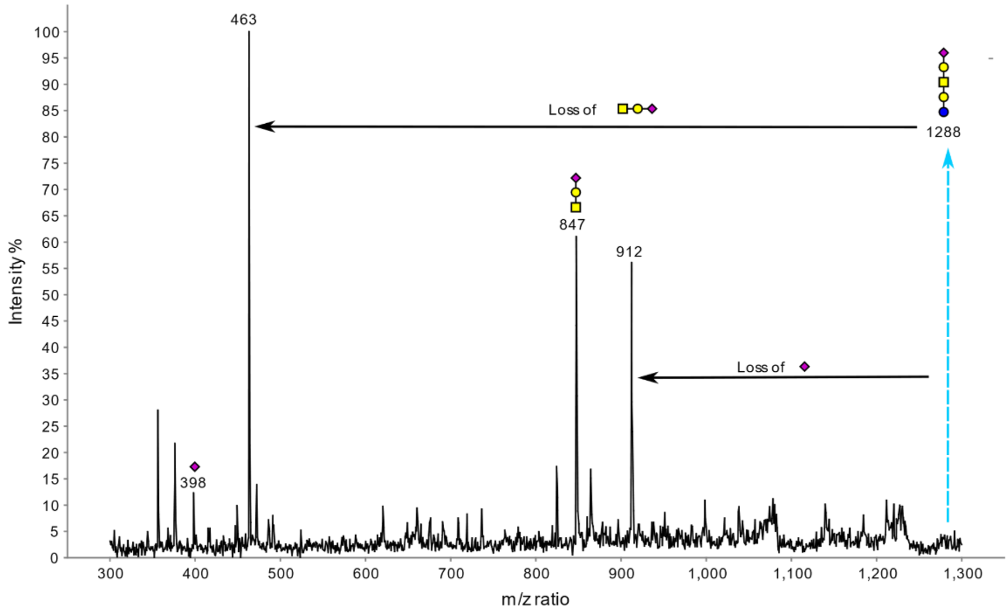


Supplementary Figure 1. MS/MS analysis of the glycolipid glycan at m/z 1288 from rested Tregs. The fragmentation of the permethylated glycan provides strong evidence that it is a GM1b.

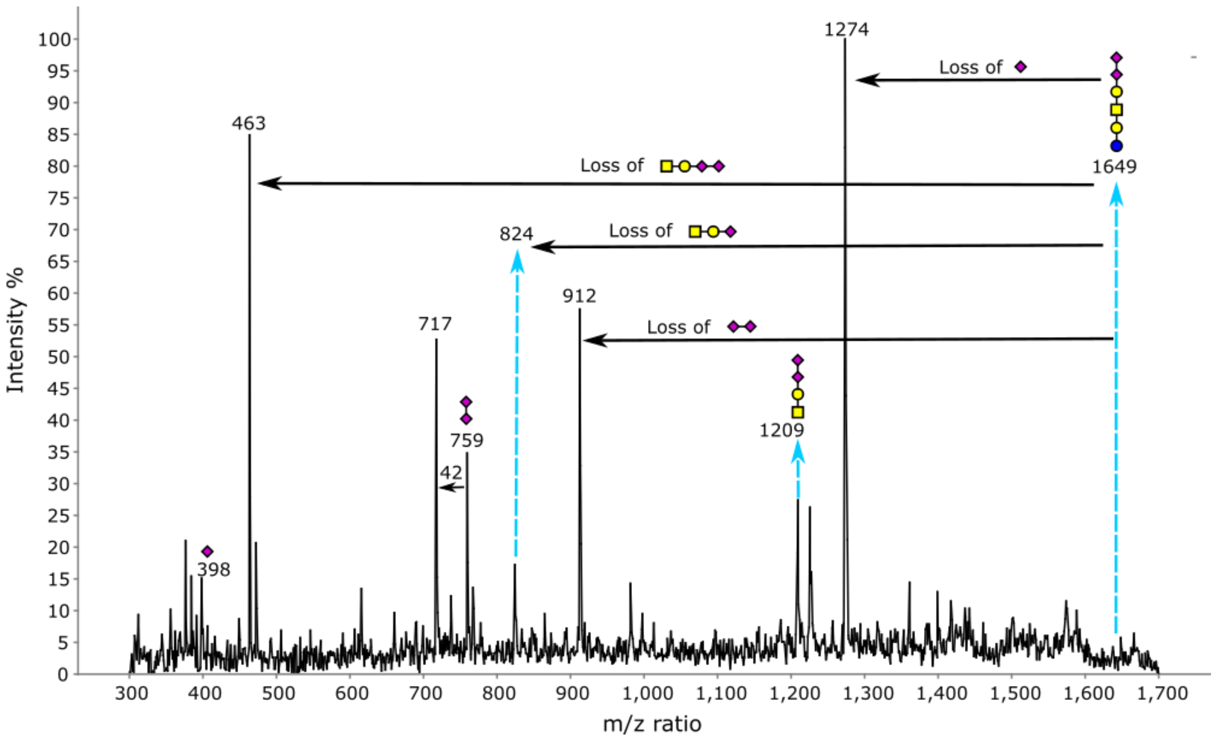


Supplementary Figure 2. MS/MS analysis of the glycolipid glycan at m/z 1649 from resting Tregs. The fragmentation of the permethylated glycan provides strong evidence that it is GD1c with two NeuAc. GD1a could coexist as a non-dominant structure. Loss of methylated carboxyl group from sialic acid was detected at m/z 717.

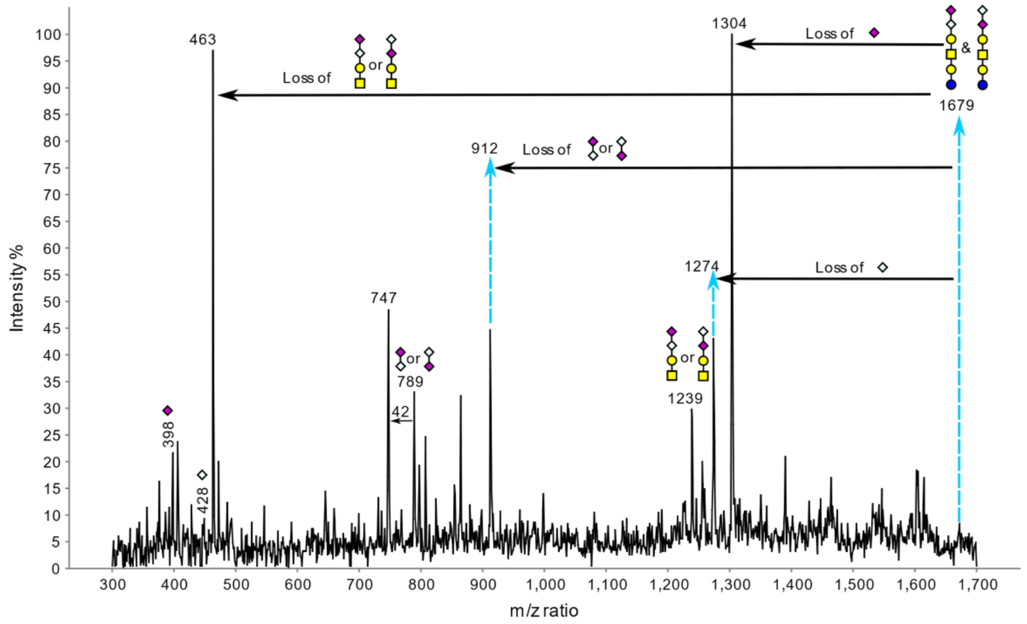


Supplementary Figure 3. MS/MS analysis of the glycolipid glycan at m/z 1679 from rested Tregs. The fragmentation of the permethylated glycan provides strong evidence that it is GD1c with 1 NeuAc and 1 NeuGc. Loss of methylated carboxyl group from sialic acid was detected at m/z 747.

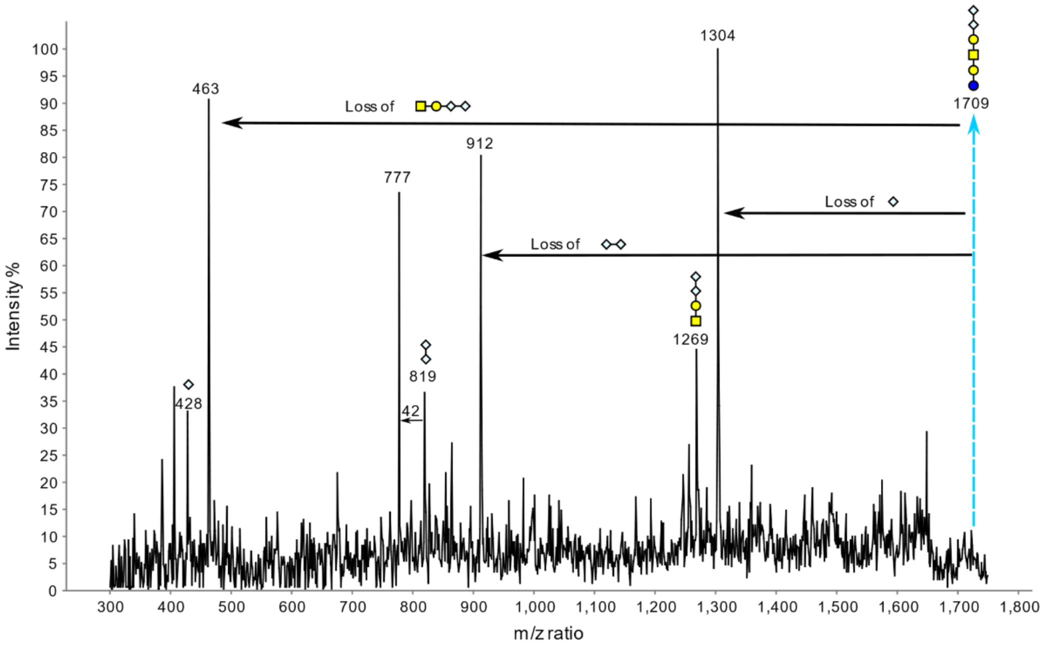


Supplementary Figure 4. MS/MS analysis of the glycolipid glycan at m/z 1709 from rested Tregs. The fragmentation of the permethylated glycan provides strong evidence that it is GD1c with 2 NeuGc. Loss of methylated carboxyl group from sialic acid was detected at m/z 777.

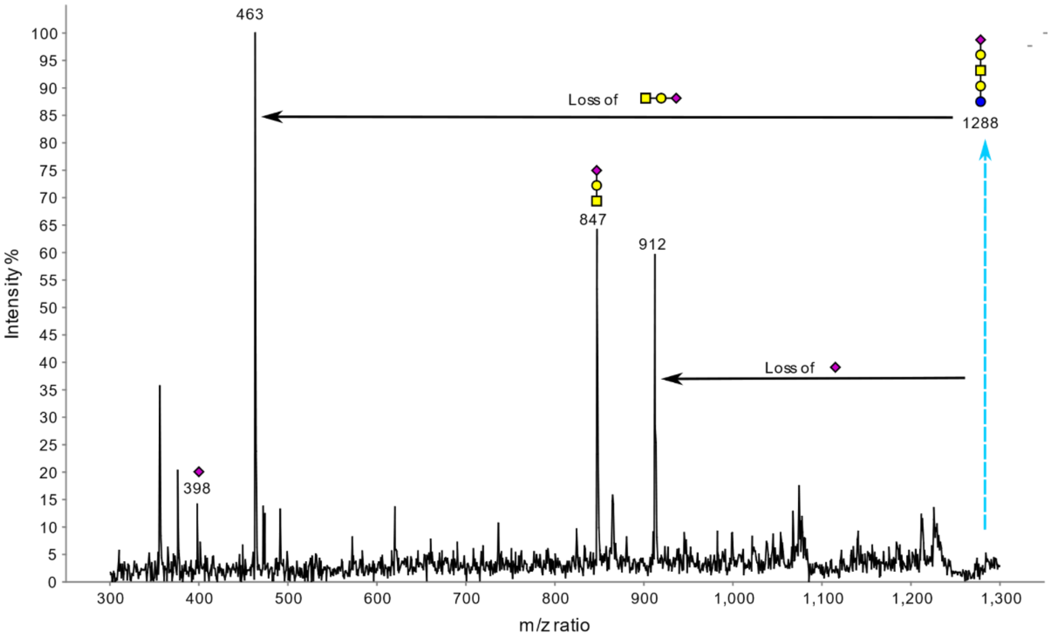


Supplementary Figure 5. MS/MS analysis of the glycolipid glycan at m/z 1288 from activated Tregs. The fragmentation of the permethylated glycan provides strong evidence that it is GM1b.

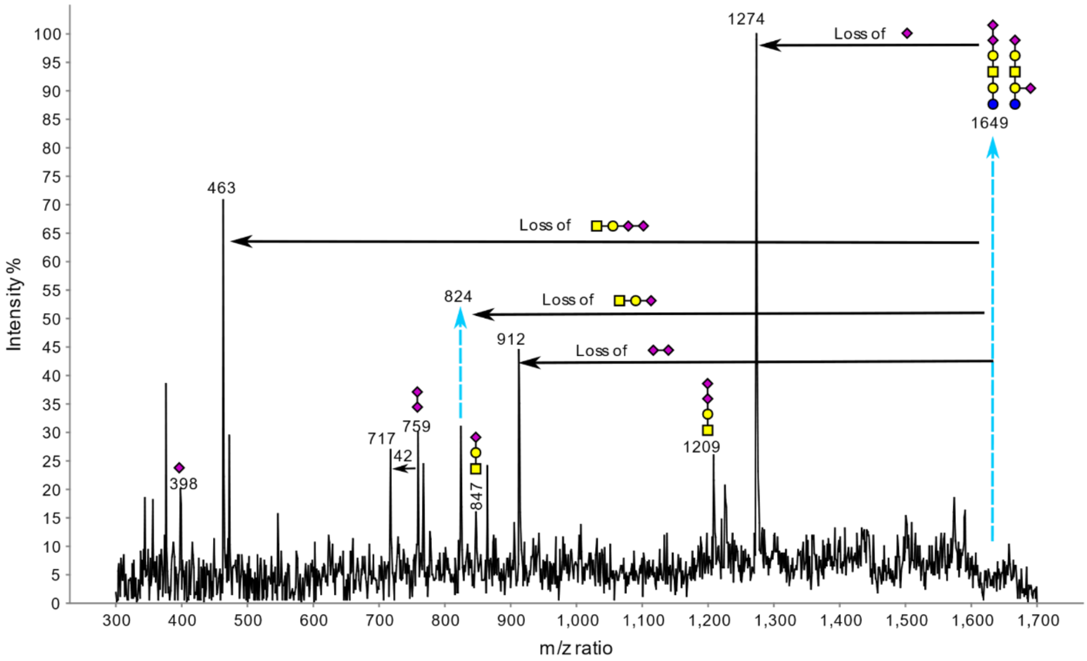


Supplementary Figure 6. MS/MS analysis of the glycolipid glycan at m/z 1649 from activated Tregs. The fragmentation of the permethylated glycan provides strong evidence that the dominant structure is GD1c with 2 NeuAc. GD1a could coexist as a non-dominant structure. Loss of methylated carboxyl group from sialic acid was detected at m/z 717.

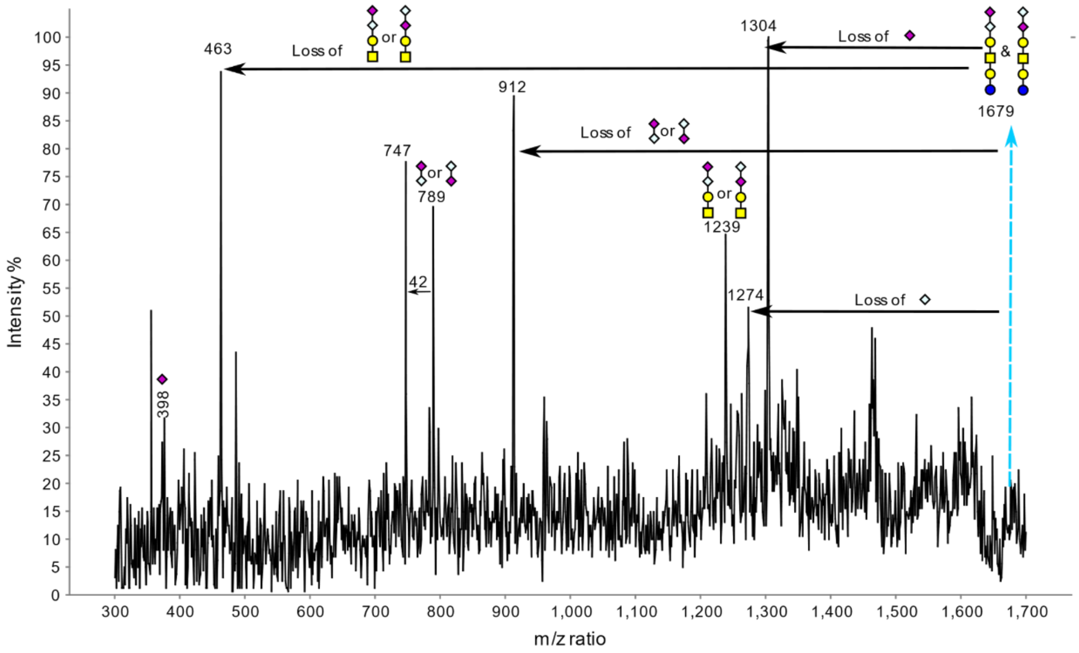


Supplementary Figure 7. MS/MS analysis of the glycolipid glycan at m/z 1679 from activated Tregs. The fragmentation of the permethylated glycan provides strong evidence that it is GD1c with 1 NeuAc and 1 NeuGc. Loss of methylated carboxyl group from sialic acid was detected at m/z 747.

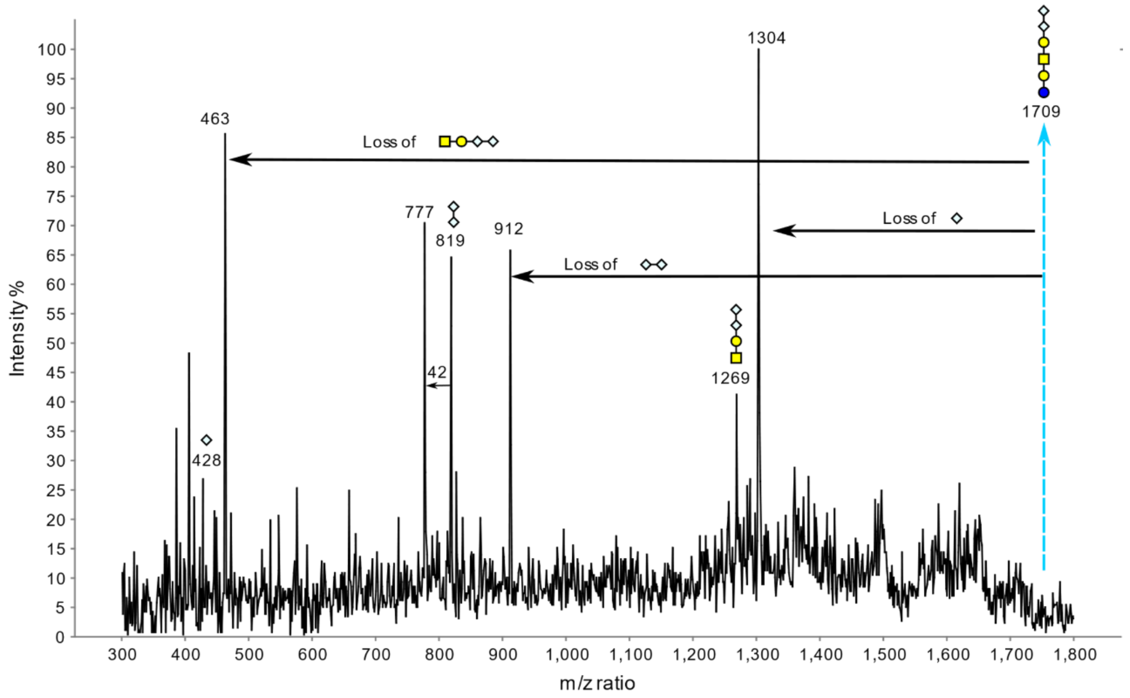


Supplementary Figure 8. MS/MS analysis of the glycolipid glycan at m/z 1709 from rest Tregs. The fragmentation of the permethylated glycan provides strong evidence that it is GD1c with 2 NeuGc. Loss of methylated carboxyl group from sialic acid was detected at m/z 777.

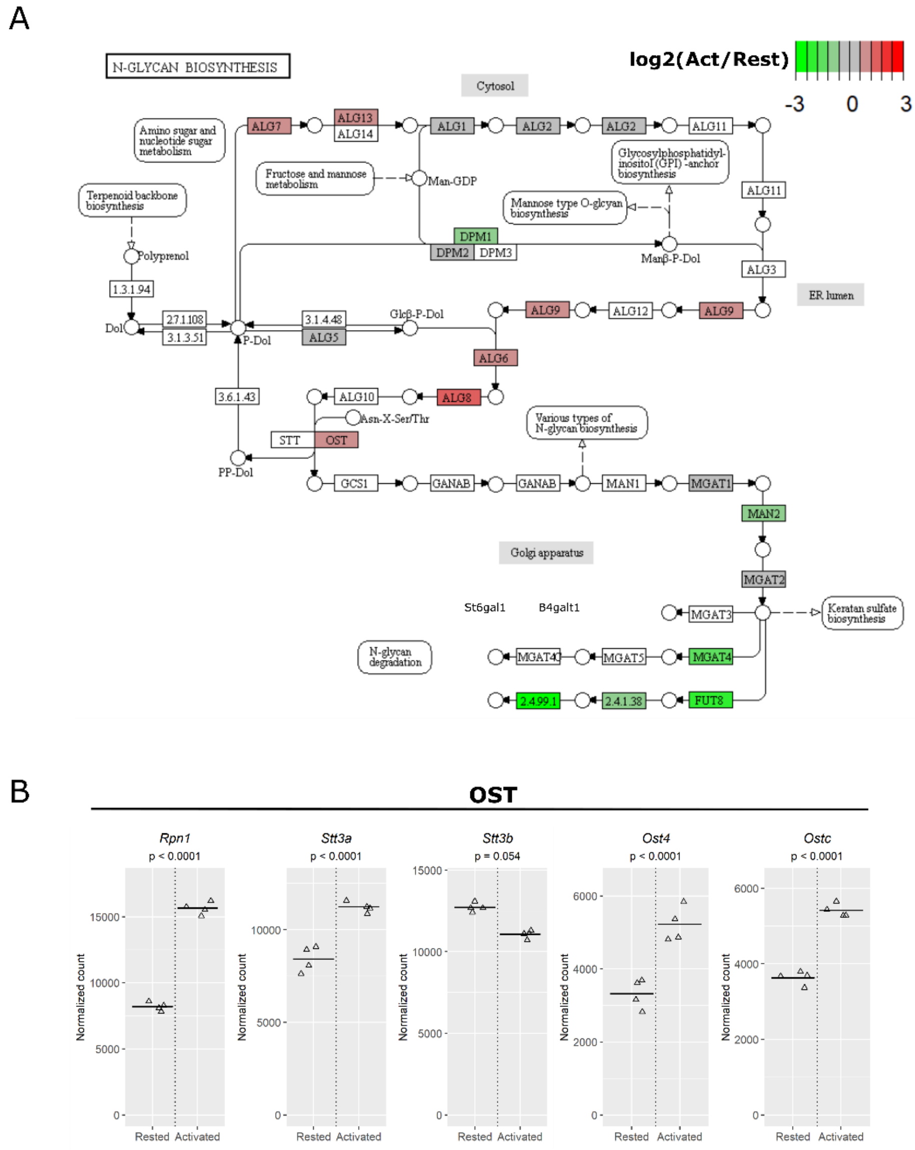


Supplementary Figure 9. Mapping RNA-Seq data to the N-glycosylation pathway. Glycosylation related genes which had normalized counts above 100 in activated cells and had significant log2 fold changes (p<0.05) were mapped to the KEGG pathway using Pathview. For the pathway node which corresponds to multiple genes, the gene which had the biggest change was used for the mapping and the change of individual genes were then manually listed. **(A)** Mapping to N-glycosylation pathway. The genes involved in synthesizing and transferring the N-glycan precursor to glycosylation sites increased upon Treg activation. **(B)** Manual listing of genes mapping to the OST node. Four of the 5 components in OST enzyme complex were increased in activated Tregs.

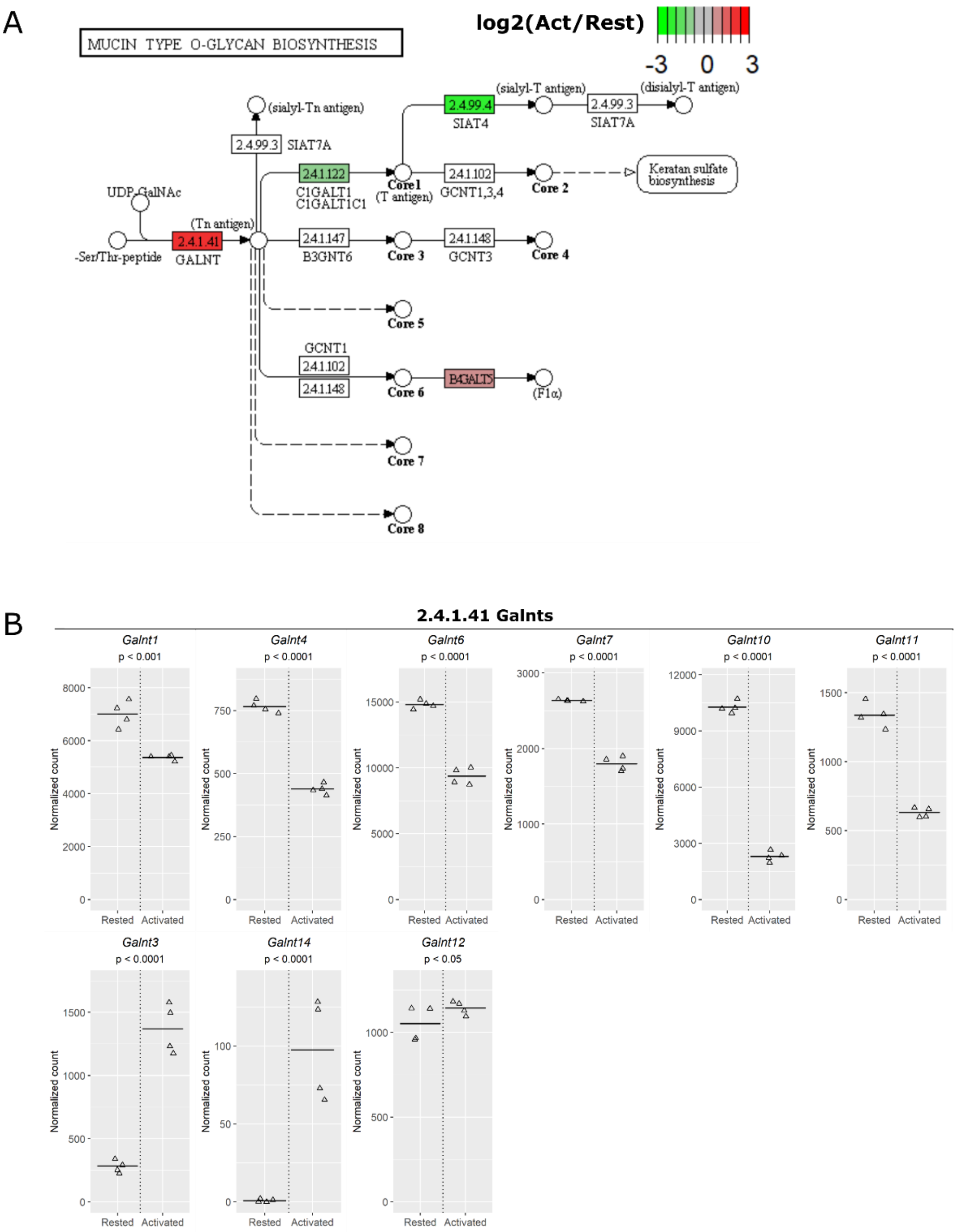


Supplementary Figure 10. Mapping RNA-Seq data to the O-glycosylation pathway. Glycosylation-related genes which had normalized counts above 100 in activated cells and significant log2 fold changes (p<0.05) were mapped to the KEGG pathway using Pathview. For the pathway node, which corresponds to multiple genes, the gene which had the biggest change was used for the mapping and the changes of individual genes were then manually listed. **(A)** Mapping to the O-glycosylation pathway. One of the *Galnts* that mapped to node 2.4.1.41 showed a large increase upon Treg activation. **(B)** Manual listing of genes that mapped to the node 2.4.1.41. *Galnt3* was the only one which showed increased expression upon Treg activation. *Galnt12* expression was unaltered and the other 6 *Galnts* showed decreased expression in activated Tregs.

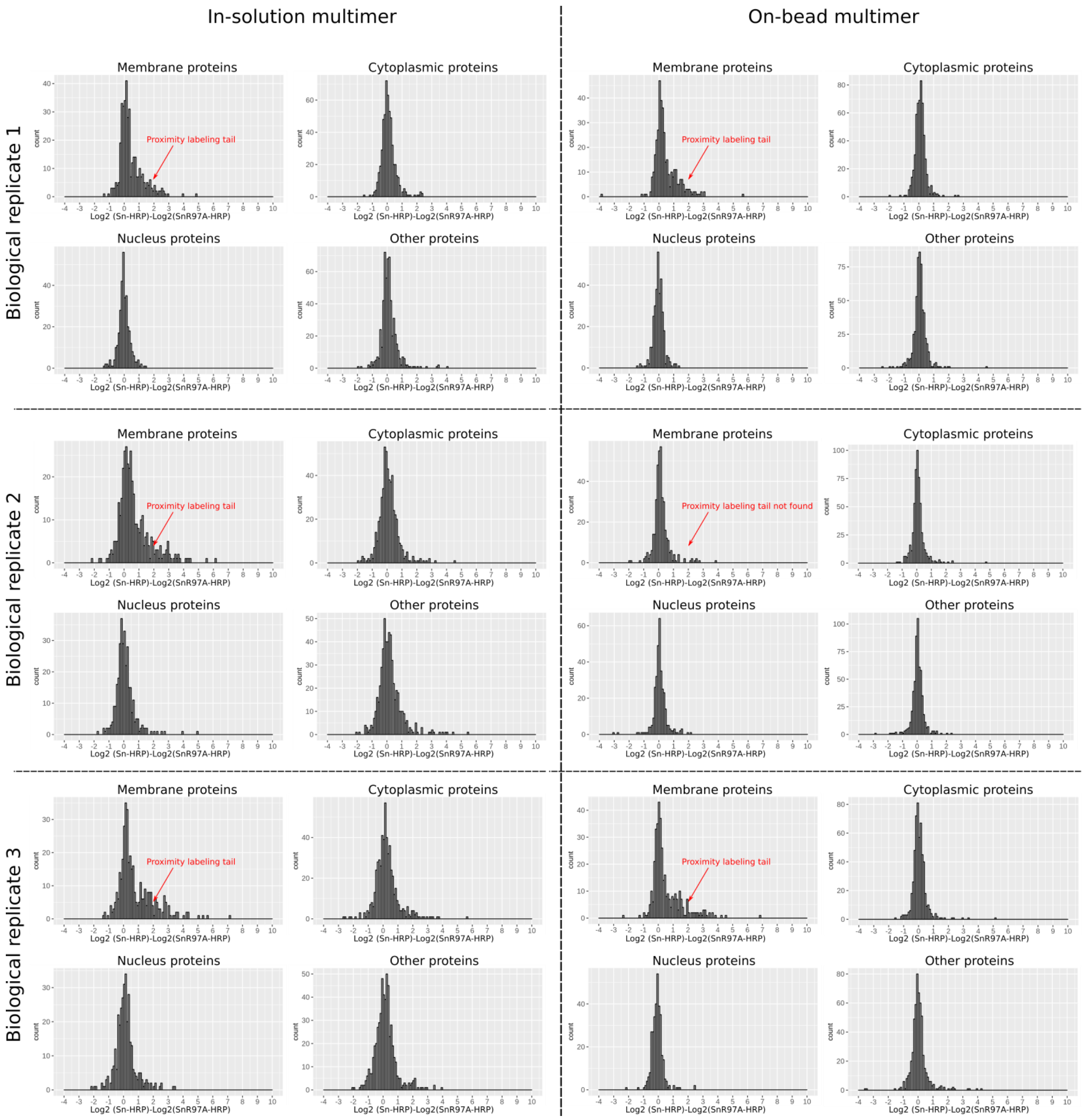


Supplementary Figure 11. Histogram of log2 fold changes of proteins in independent proximity labelling experiments and their negative controls. The proximity labelling tail was found in 5 of the 6 proximity labelling experiments as indicated.

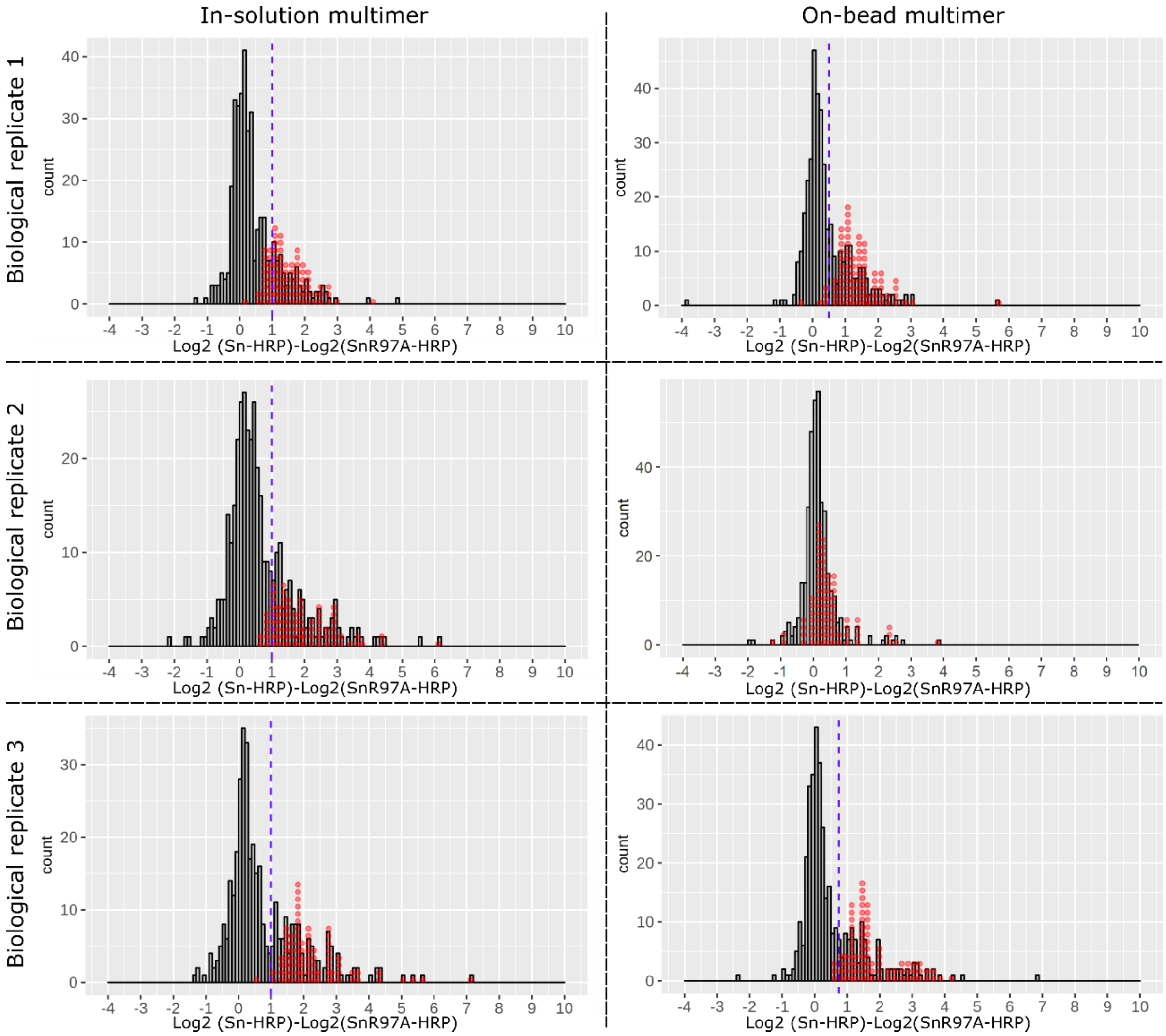


Supplementary Figure 12. The glycosylated proteins from the significant hits on the volcano plot were mapped back to the individual histogram of total membrane proteins. Each red dot on the histograms represents a significant hit on the volcano plot. The red dots outside the proximity labelling tail (on the left side of the blue doted vertical line in each histogram) were filtered out. The on-bead multimer proximity labelling of the second biological replicate did not have the proximity labelling tail and was not used for data filtration.

Supplementary Table 1. List of glycosylation-related genes and their expression levels in Tregs

| Gene ID | Gene name | log2(Act/Rest) | p value | Average normalized count Rest | Average normalized count Act |
|---|---|---|---|---|---|
| ENSMUSG00000024064 | Galnt14 | 6.899406382 | 1.80E-10 | 0.7 | 97.525 |
| ENSMUSG00000050473 | Slc35d3 | 5.738498467 | 2.82E-08 | 0.45 | 31.325 |
| ENSMUSG00000094651 | Gal3st2 | 4.915428974 | 0.00155259 | 0 | 3.575 |
| ENSMUSG00000033350 | Chst2 | 4.104136643 | 3.74E-12 | 68.375 | 1111.225 |
| ENSMUSG00000042684 | Npl | 3.414816705 | 5.10E-07 | 2.45 | 26.025 |
| ENSMUSG00000021194 | Chga | 2.827898966 | 0.000364741 | 0.975 | 7.425 |
| ENSMUSG00000006800 | Sulf2 | 2.453983682 | 2.22E-12 | 243.525 | 1269.125 |
| ENSMUSG00000026994 | Galnt3 | 2.34341527 | 6.59E-12 | 284.975 | 1369.675 |
| ENSMUSG00000079442 | St6galnac4 | 2.329156188 | 7.35E-17 | 1608.975 | 7676.7 |
| ENSMUSG00000034780 | B3galt1 | 2.124359596 | 0.011350253 | 1.625 | 6.625 |
| ENSMUSG00000042428 | Mgat3 | 2.053080864 | 0.003242152 | 1.875 | 8.1 |
| ENSMUSG00000028541 | B4galt2 | 2.029300447 | 4.18E-06 | 23.65 | 91.45 |
| ENSMUSG00000026811 | St6galnac6 | 1.842531026 | 3.60E-11 | 452.3 | 1539.9 |
| ENSMUSG00000032854 | Ugt8a | 1.629497132 | 0.280524412 | 0 | 0.275 |
| ENSMUSG00000034040 | Galnt17 | 1.629497132 | 0.280524412 | 0 | 0.275 |
| ENSMUSG00000047658 | Gal3st3 | 1.629497132 | 0.280524412 | 0 | 0.275 |
| ENSMUSG00000047878 | A4galt | 1.629497132 | 0.280524412 | 0 | 0.275 |
| ENSMUSG00000021130 | Galnt16 | 1.624693106 | 0.277888517 | 0 | 0.25 |
| ENSMUSG00000090145 | Ugt1a6b | 1.624693106 | 0.277888517 | 0 | 0.25 |
| ENSMUSG00000090165 | Ugt1a10 | 1.624693106 | 0.277888517 | 0 | 0.25 |
| ENSMUSG00000034118 | Tpst1 | 1.622282144 | 6.65E-09 | 255.425 | 748.475 |
| ENSMUSG00000040434 | Large2 | 1.563485295 | 0.001018153 | 5.375 | 15.425 |
| ENSMUSG00000035704 | Alg8 | 1.549310569 | 4.10E-13 | 1049.625 | 2914 |
| ENSMUSG00000031387 | Renbp | 1.525152185 | 6.00E-08 | 91.675 | 251.45 |
| ENSMUSG00000027221 | Chst1 | 1.387403794 | 0.147864061 | 0.75 | 2.025 |
| ENSMUSG00000020766 | Galk1 | 1.361698307 | 8.75E-13 | 2549.575 | 6218.8 |
| ENSMUSG00000058152 | Chsy3 | 1.310745503 | 0.334019854 | 0.225 | 0.725 |
| ENSMUSG00000061731 | Ext1 | 1.306565843 | 2.24E-08 | 733.775 | 1719.225 |
| ENSMUSG00000048807 | Slc35e4 | 1.252323391 | 0.000106466 | 47.15 | 106.625 |
| ENSMUSG00000026156 | B3gat2 | 1.235704984 | 0.059532918 | 1.925 | 4.55 |
| ENSMUSG00000010047 | Hyal2 | 1.232925335 | 1.16E-08 | 380.7 | 849.325 |
| ENSMUSG00000056091 | St3gal5 | 1.159553278 | 9.42E-06 | 50.95 | 108.325 |
| ENSMUSG00000031266 | Gla | 1.15599329 | 1.54E-07 | 243.15 | 514.05 |
| ENSMUSG00000032123 | Dpagt1 | 1.066058046 | 1.57E-10 | 1090.575 | 2166.375 |
| ENSMUSG00000034612 | Chst11 | 1.064687612 | 1.66E-05 | 330.975 | 655.175 |
| ENSMUSG00000032059 | Alg9 | 1.010454071 | 6.69E-10 | 844.4 | 1616.325 |
| ENSMUSG00000030062 | Rpn1 | 1.006517452 | 3.18E-11 | 8207.75 | 15652.475 |
| ENSMUSG00000021978 | Extl3 | 1.006405328 | 5.67E-10 | 3787.35 | 7221.2 |
| ENSMUSG00000041718 | Alg13 | 1.003045631 | 9.01E-08 | 464.45 | 884.375 |
| ENSMUSG00000026670 | Uap1 | 0.948766175 | 3.29E-11 | 3192.275 | 5852.975 |

| | | | | | |
|---|---|---|---|---|---|
| ENSMUSG00000022174 | Dad1 | 0.859903207 | 7.32E-08 | 4413.7 | 7592.8 |
| ENSMUSG00000028334 | Nans | 0.841082221 | 5.59E-08 | 4551 | 7730.5 |
| ENSMUSG00000027977 | Ndst3 | 0.818592186 | 0.551305324 | 0.275 | 0.5 |
| ENSMUSG00000028671 | Gale | 0.818214826 | 1.39E-06 | 902.775 | 1510.425 |
| ENSMUSG00000017929 | B4galt5 | 0.814145236 | 3.05E-06 | 2849.625 | 4749.4 |
| ENSMUSG00000026342 | Slc35f5 | 0.775657335 | 1.38E-05 | 227.75 | 371.2 |
| ENSMUSG00000028521 | Slc35d1 | 0.764480563 | 5.32E-08 | 771.125 | 1243.8 |
| ENSMUSG00000070284 | Gmppb | 0.757953642 | 2.27E-08 | 1625.575 | 2609.4 |
| ENSMUSG00000032306 | Mpi | 0.750445665 | 1.08E-05 | 487.925 | 779.55 |
| ENSMUSG00000073792 | Alg6 | 0.736775488 | 2.75E-05 | 229.975 | 363.85 |
| ENSMUSG00000031803 | B3gnt3 | 0.694263196 | 0.00024811 | 164.8 | 253.925 |
| ENSMUSG00000042195 | Slc35f2 | 0.685655266 | 5.17E-06 | 496.75 | 758.2 |
| ENSMUSG00000039254 | Pomt1 | 0.679105257 | 2.03E-05 | 443.225 | 673.375 |
| ENSMUSG00000041372 | B4galnt3 | 0.676016095 | 0.223288663 | 2.95 | 4.525 |
| ENSMUSG00000028757 | Ddost | 0.664808452 | 5.14E-09 | 7457.7 | 11229.125 |
| ENSMUSG00000020873 | Slc35b1 | 0.626743649 | 7.86E-07 | 2683.475 | 3928.925 |
| ENSMUSG00000034744 | Nagk | 0.621129547 | 4.98E-05 | 525.35 | 766.375 |
| ENSMUSG00000047379 | B4gat1 | 0.602367841 | 0.218109324 | 5.325 | 7.8 |
| ENSMUSG00000003418 | St8sia6 | 0.598633784 | 0.001954882 | 338.275 | 485.825 |
| ENSMUSG00000037347 | Chst7 | 0.580004958 | 0.325222354 | 7.675 | 10.975 |
| ENSMUSG00000007038 | Neu1 | 0.552888056 | 2.45E-06 | 1068 | 1487 |
| ENSMUSG00000036632 | Alg5 | 0.545654931 | 7.59E-07 | 2187.625 | 3031.85 |
| ENSMUSG00000029992 | Gfpt1 | 0.507083473 | 2.88E-06 | 1911.875 | 2579.8 |
| ENSMUSG00000025534 | Gusb | 0.476362442 | 7.50E-06 | 2334.975 | 3088.625 |
| ENSMUSG00000036587 | Fut7 | 0.45838241 | 7.74E-05 | 2878.825 | 3757.2 |
| ENSMUSG00000022570 | Tsta3 | 0.389225541 | 2.69E-05 | 2233.475 | 2777.375 |
| ENSMUSG00000027642 | Rpn2 | 0.383958888 | 1.42E-05 | 12187.775 | 15107.975 |
| ENSMUSG00000037722 | Gnpnat1 | 0.380848245 | 0.000557993 | 1153.3 | 1425.925 |
| ENSMUSG00000017664 | Slc35c2 | 0.377641568 | 0.002207688 | 1700.85 | 2093.7 |
| ENSMUSG00000057363 | Uxs1 | 0.373023669 | 0.000368477 | 1505.05 | 1853.875 |
| ENSMUSG00000038372 | Gmds | 0.363482019 | 0.000740956 | 959 | 1170.725 |
| ENSMUSG00000035778 | Ggta1 | 0.358731519 | 0.001766785 | 2083.325 | 2543 |
| ENSMUSG00000001891 | Ugp2 | 0.356229289 | 0.000111977 | 2417.65 | 2940.675 |
| ENSMUSG00000037089 | Slc35b2 | 0.347880484 | 9.17E-05 | 1941.925 | 2348.5 |
| ENSMUSG00000028700 | Pomgnt1 | 0.341950181 | 0.000136417 | 1731.05 | 2083.525 |
| ENSMUSG00000034126 | Pomt2 | 0.326302441 | 0.003128145 | 529.95 | 630.775 |
| ENSMUSG00000079469 | Pigb | 0.303270926 | 0.001643682 | 982.25 | 1150.925 |
| ENSMUSG00000048970 | C1galt1c1 | 0.30052855 | 0.013722453 | 557.15 | 652 |
| ENSMUSG00000020346 | Mgat1 | 0.298649697 | 0.000439991 | 5198.175 | 6075.75 |
| ENSMUSG00000026810 | Dpm2 | 0.293342847 | 0.002646122 | 1844.5 | 2144.475 |
| ENSMUSG00000038702 | Dsel | 0.284673794 | 0.093588544 | 235.125 | 272.125 |
| ENSMUSG00000030930 | Chst15 | 0.263546738 | 0.122542951 | 102.4 | 116.85 |
| ENSMUSG00000052423 | B4galt3 | 0.253260071 | 0.004365317 | 1461.8 | 1654.05 |
| ENSMUSG00000031156 | Slc35a2 | 0.197831392 | 0.043429874 | 871.975 | 948.875 |
| ENSMUSG00000039774 | Galnt12 | 0.196365221 | 0.035958555 | 1051.6 | 1143.75 |

| | | | | | |
|---|---|---|---|---|---|
| ENSMUSG00000032997 | Chpf | 0.193319806 | 0.031280498 | 1841.65 | 1996.95 |
| ENSMUSG00000039427 | Alg1 | 0.192105187 | 0.021351211 | 1233.8 | 1338.45 |
| ENSMUSG00000020363 | Gfpt2 | 0.168247575 | 0.421181986 | 232.375 | 247.475 |
| ENSMUSG00000031521 | Aga | 0.16411405 | 0.097448736 | 1009.425 | 1074.6 |
| ENSMUSG00000015027 | Galns | 0.151779966 | 0.114321765 | 1013.75 | 1068.925 |
| ENSMUSG00000043300 | B3galnt1 | 0.151106796 | 0.620415574 | 48.4 | 51.025 |
| ENSMUSG00000025158 | Rfng | 0.145933991 | 0.122906778 | 892.125 | 937.175 |
| ENSMUSG00000016756 | Cmah | 0.129529131 | 0.629169998 | 243.65 | 254.075 |
| ENSMUSG00000035845 | Alg12 | 0.123871545 | 0.189501449 | 803.25 | 830.875 |
| ENSMUSG00000036427 | Gpi1 | 0.107707215 | 0.113988 | 21655.525 | 22165.525 |
| ENSMUSG00000039628 | Hs3st6 | 0.107568216 | 0.924034102 | 0.725 | 0.775 |
| ENSMUSG00000025728 | Pigq | 0.106805793 | 0.130639439 | 3050.025 | 3116.85 |
| ENSMUSG00000045594 | Glb1 | 0.105690546 | 0.211602767 | 2387.6 | 2437.425 |
| ENSMUSG00000050796 | B3galt6 | 0.100945511 | 0.340723 | 456.2 | 464.375 |
| ENSMUSG00000062184 | Hs6st2 | 0.099670421 | 0.910136285 | 1.825 | 1.8 |
| ENSMUSG00000028538 | St3gal3 | 0.089006526 | 0.314208134 | 810.65 | 818.625 |
| ENSMUSG00000022367 | Has2 | 0.081191182 | 0.95595038 | 0.225 | 0.25 |
| ENSMUSG00000035239 | Neu3 | 0.080218946 | 0.653769454 | 389.3 | 391.85 |
| ENSMUSG00000025791 | Pgm1 | 0.0762477 | 0.360410775 | 1844.775 | 1849.8 |
| ENSMUSG00000036155 | Mgat5 | 0.070946832 | 0.387736348 | 3124.1 | 3112.45 |
| ENSMUSG00000060181 | Slc35e3 | 0.068043562 | 0.618550487 | 285.325 | 283.75 |
| ENSMUSG00000029201 | Ugdh | 0.0645079 | 0.368002176 | 2305.55 | 2292.175 |
| ENSMUSG00000031753 | Cog4 | 0.04257882 | 0.516273673 | 2248.375 | 2198.95 |
| ENSMUSG00000046020 | Pofut1 | 0.023170121 | 0.803154481 | 1331.725 | 1286.075 |
| ENSMUSG00000032252 | Glce | 0.018786941 | 0.859178871 | 583.55 | 562.075 |
| ENSMUSG00000027207 | Galk2 | 0.005165825 | 0.94597718 | 1904.25 | 1814.975 |
| ENSMUSG00000033021 | Gmppa | -0.006007477 | 0.93792462 | 2474.525 | 2338.175 |
| ENSMUSG00000032038 | St3gal4 | -0.021152088 | 0.752091409 | 2570.6 | 2404.25 |
| ENSMUSG00000020260 | Pofut2 | -0.02389197 | 0.746377076 | 1663.95 | 1553.65 |
| ENSMUSG00000056131 | Pgm3 | -0.025117827 | 0.78480519 | 794.675 | 741.175 |
| ENSMUSG00000071649 | B3gat3 | -0.038575082 | 0.670876222 | 2524.05 | 2330.5 |
| ENSMUSG00000033272 | Slc35a4 | -0.047129814 | 0.452048554 | 4825.125 | 4434.525 |
| ENSMUSG00000017760 | Ctsa | -0.056562939 | 0.371620421 | 5739.6 | 5237.6 |
| ENSMUSG00000049307 | Fut4 | -0.057668229 | 0.663943724 | 362.425 | 331.525 |
| ENSMUSG00000027957 | Slc35a3 | -0.06659395 | 0.48644941 | 2320.175 | 2105.15 |
| ENSMUSG00000022474 | Pmm1 | -0.072505038 | 0.652481609 | 914.9 | 823.5 |
| ENSMUSG00000036599 | Chst12 | -0.087117638 | 0.359505556 | 2456.125 | 2193.625 |
| ENSMUSG00000022793 | B4galt4 | -0.108285676 | 0.384285267 | 796.6 | 703.65 |
| ENSMUSG00000037470 | Uggt1 | -0.108593218 | 0.081213694 | 7403.275 | 6517.825 |
| ENSMUSG00000006731 | B4galnt1 | -0.115630014 | 0.06849394 | 5998.225 | 5256.475 |
| ENSMUSG00000069920 | B3gnt9 | -0.121747582 | 0.528634359 | 96.6 | 84.125 |
| ENSMUSG00000022940 | Pigp | -0.122418219 | 0.22563163 | 977.7 | 854.025 |
| ENSMUSG00000038843 | Gcnt1 | -0.128544175 | 0.529821108 | 985.75 | 853 |
| ENSMUSG00000030282 | Cmas | -0.136776425 | 0.051057335 | 3269.4 | 2824.8 |
| ENSMUSG00000028838 | Extl1 | -0.158155317 | 0.373007301 | 173.75 | 147.8 |

| | | | | | |
|---|---|---|---|---|---|
| ENSMUSG00000075470 | Alg10b | -0.161219105 | 0.071032282 | 2486.6 | 2113.375 |
| ENSMUSG00000043998 | Mgat2 | -0.162985026 | 0.023825216 | 3715.225 | 3152.35 |
| ENSMUSG00000004383 | Large1 | -0.170369803 | 0.368305538 | 191.675 | 161.6 |
| ENSMUSG00000031979 | Cog2 | -0.172744226 | 0.03891711 | 2416.525 | 2036.075 |
| ENSMUSG00000031591 | Asah1 | -0.182814791 | 0.032977316 | 1645.15 | 1377.325 |
| ENSMUSG00000028048 | Gba | -0.18614027 | 0.016380473 | 2854.175 | 2380.2 |
| ENSMUSG00000019731 | Slc35e1 | -0.200355728 | 0.012595297 | 2506.8 | 2070.85 |
| ENSMUSG00000025232 | Hexa | -0.265591823 | 0.002201037 | 1642.125 | 1297.475 |
| ENSMUSG00000034893 | Cog3 | -0.276048882 | 0.001021647 | 3259.35 | 2558.775 |
| ENSMUSG00000039357 | Fut11 | -0.278026401 | 0.00812758 | 686.55 | 537.6 |
| ENSMUSG00000027198 | Ext2 | -0.290088218 | 0.000470154 | 3199.05 | 2484.6 |
| ENSMUSG00000016534 | Lamp2 | -0.308246614 | 0.002429299 | 3206.425 | 2460.6 |
| ENSMUSG00000000420 | Galnt1 | -0.311616346 | 0.000211587 | 7010.275 | 5365.125 |
| ENSMUSG00000045216 | Hs6st1 | -0.316620193 | 0.002142148 | 1146.075 | 874.5 |
| ENSMUSG00000022711 | Pmm2 | -0.327861275 | 7.55E-05 | 3326.25 | 2515.8 |
| ENSMUSG00000032640 | Chsy1 | -0.335641429 | 0.000956731 | 3463.9 | 2601.6 |
| ENSMUSG00000027742 | Cog6 | -0.352138875 | 0.00017028 | 1907.95 | 1420.65 |
| ENSMUSG00000010051 | Hyal1 | -0.353616635 | 0.264595038 | 23.075 | 17.275 |
| ENSMUSG00000028479 | Gne | -0.36807707 | 5.97E-05 | 2388.325 | 1758.95 |
| ENSMUSG00000031916 | Cog8 | -0.368305913 | 4.94E-05 | 2297.65 | 1689.8 |
| ENSMUSG00000032226 | Gcnt3 | -0.390837946 | 0.751611412 | 0.75 | 0.5 |
| ENSMUSG00000039308 | Ndst2 | -0.393698051 | 0.000240162 | 1494.3 | 1080.125 |
| ENSMUSG00000039740 | Alg2 | -0.394978606 | 0.001619021 | 999.525 | 722.325 |
| ENSMUSG00000018169 | Mfng | -0.395138716 | 1.84E-05 | 5933.5 | 4289.875 |
| ENSMUSG00000057060 | Slc35f3 | -0.396003262 | 0.751318652 | 0.775 | 0.5 |
| ENSMUSG00000021360 | Gcnt2 | -0.397680815 | 0.026024235 | 680.6 | 489.3 |
| ENSMUSG00000022453 | Naga | -0.399746206 | 9.72E-05 | 1476.675 | 1062.85 |
| ENSMUSG00000038886 | Man2a2 | -0.406130561 | 0.000464352 | 2667.65 | 1912.2 |
| ENSMUSG00000021003 | Galc | -0.410704107 | 0.0061298 | 280.55 | 200.025 |
| ENSMUSG00000074916 | Chst14 | -0.413065408 | 0.003911335 | 426.275 | 303.75 |
| ENSMUSG00000040151 | Hs2st1 | -0.417445605 | 0.000364016 | 831.075 | 590.9 |
| ENSMUSG00000020868 | Xylt2 | -0.425765064 | 6.06E-05 | 1452.5 | 1026.95 |
| ENSMUSG00000036620 | Mgat4b | -0.429127356 | 0.000976764 | 611.025 | 430.775 |
| ENSMUSG00000005142 | Man2b1 | -0.447801759 | 7.39E-07 | 8817.725 | 6138.6 |
| ENSMUSG00000018661 | Cog1 | -0.458705536 | 2.38E-06 | 2667.65 | 1844.125 |
| ENSMUSG00000025579 | Gaa | -0.469871926 | 0.000143004 | 1635.475 | 1119.55 |
| ENSMUSG00000039242 | B3galnt2 | -0.474335624 | 1.84E-05 | 1989.175 | 1358.775 |
| ENSMUSG00000031608 | Galnt7 | -0.477911922 | 1.69E-06 | 2638.55 | 1800.175 |
| ENSMUSG00000028293 | Slc35a1 | -0.480518353 | 1.43E-05 | 2124.975 | 1444.85 |
| ENSMUSG00000015189 | Casd1 | -0.482162047 | 1.43E-05 | 2002.55 | 1362.825 |
| ENSMUSG00000036073 | Galt | -0.482981258 | 0.00017214 | 1269.95 | 861.575 |
| ENSMUSG00000022686 | B3gnt5 | -0.48764857 | 0.013947578 | 361.225 | 244.75 |
| ENSMUSG00000029344 | Tpst2 | -0.496679061 | 8.40E-06 | 2483.975 | 1672.2 |
| ENSMUSG00000031447 | Lamp1 | -0.500446089 | 2.66E-05 | 10937.325 | 7353 |
| ENSMUSG00000055629 | B4galnt4 | -0.501145649 | 0.101488169 | 1775.325 | 1183.175 |

| | | | | | |
|---|---|---|---|---|---|
| ENSMUSG00000037012 | Hk1 | -0.502378479 | 4.94E-06 | 11475.7 | 7707.125 |
| ENSMUSG00000047712 | Ust | -0.531515879 | 0.000311953 | 478.375 | 314.45 |
| ENSMUSG00000042042 | Csgalnact2 | -0.542489472 | 4.50E-05 | 842.175 | 549.275 |
| ENSMUSG00000033316 | Galnt9 | -0.580007841 | 0.07630275 | 70.95 | 45.175 |
| ENSMUSG00000037280 | Galnt6 | -0.585466456 | 8.27E-08 | 14817.775 | 9385.2 |
| ENSMUSG00000028413 | B4galt1 | -0.603867086 | 1.52E-05 | 25161.575 | 15737.725 |
| ENSMUSG00000029209 | Gnpda2 | -0.610008624 | 3.11E-05 | 1054.65 | 656.85 |
| ENSMUSG00000037049 | Smpd1 | -0.613639956 | 2.42E-06 | 2042.475 | 1266.1 |
| ENSMUSG00000000594 | Gm2a | -0.615204354 | 3.38E-05 | 10122.875 | 6263.875 |
| ENSMUSG00000050229 | Pigm | -0.629312599 | 1.21E-07 | 2717.725 | 1667.025 |
| ENSMUSG00000041731 | Pgm5 | -0.649352621 | 0.63988336 | 0.525 | 0.275 |
| ENSMUSG00000033114 | Slc35d2 | -0.649720377 | 2.29E-05 | 548.6 | 331.825 |
| ENSMUSG00000024085 | Man2a1 | -0.65630159 | 8.41E-08 | 3456.525 | 2082.925 |
| ENSMUSG00000025220 | Mgea5 | -0.657106071 | 1.13E-07 | 19498.125 | 11747.625 |
| ENSMUSG00000042460 | C1galt1 | -0.667627385 | 3.03E-08 | 2617.025 | 1565.075 |
| ENSMUSG00000052102 | Gnpda1 | -0.670497953 | 1.74E-06 | 2216.625 | 1320.975 |
| ENSMUSG00000053870 | Fpgt | -0.677814703 | 1.59E-05 | 911.775 | 541.625 |
| ENSMUSG00000049922 | Slc35c1 | -0.684809022 | 1.30E-06 | 2392.9 | 1416.45 |
| ENSMUSG00000001751 | Naglu | -0.697206106 | 1.56E-06 | 993.475 | 582.5 |
| ENSMUSG00000001942 | Siae | -0.716505903 | 7.09E-07 | 856.725 | 495.5 |
| ENSMUSG00000029171 | Pgm2 | -0.7197686 | 1.33E-07 | 1395.225 | 804.65 |
| ENSMUSG00000090035 | Galnt4 | -0.726371741 | 7.13E-07 | 765.45 | 439.2 |
| ENSMUSG00000034951 | Cog7 | -0.72672344 | 6.93E-07 | 999.475 | 573.775 |
| ENSMUSG00000027963 | Extl2 | -0.738795665 | 0.000303762 | 527.725 | 300.425 |
| ENSMUSG00000031381 | Piga | -0.755461947 | 7.75E-09 | 2203.175 | 1239.25 |
| ENSMUSG00000022620 | Arsa | -0.761080009 | 4.58E-07 | 800.925 | 449.2 |
| ENSMUSG00000030729 | Pgm2l1 | -0.791855835 | 5.68E-07 | 719.45 | 394.3 |
| ENSMUSG00000034160 | Ogt | -0.792307971 | 1.54E-09 | 19940.825 | 10932.65 |
| ENSMUSG00000051022 | Hs3st1 | -0.798977693 | 0.002407959 | 102.075 | 55.95 |
| ENSMUSG00000026956 | Uap1l1 | -0.815144554 | 1.11E-07 | 1055.6 | 570.05 |
| ENSMUSG00000021432 | Slc35b3 | -0.817026506 | 6.77E-08 | 3263.475 | 1760.625 |
| ENSMUSG00000022664 | Slc35a5 | -0.819058825 | 1.02E-05 | 1050.425 | 566.6 |
| ENSMUSG00000078919 | Dpm1 | -0.844126819 | 1.74E-05 | 247.725 | 131.025 |
| ENSMUSG00000033540 | Idua | -0.853437496 | 2.35E-07 | 731.35 | 384.8 |
| ENSMUSG00000029570 | Lfng | -0.855895687 | 7.86E-09 | 15674.575 | 8233.075 |
| ENSMUSG00000070407 | Hs3st3b1 | -0.886127892 | 3.60E-06 | 409.9 | 210.525 |
| ENSMUSG00000028381 | Ugcg | -0.954580502 | 6.75E-11 | 8395.85 | 4113.725 |
| ENSMUSG00000024781 | Lipa | -0.958222138 | 1.02E-09 | 1572.35 | 769 |
| ENSMUSG00000028164 | Manba | -0.959038814 | 1.86E-09 | 1433.325 | 699.475 |
| ENSMUSG00000028032 | Papss1 | -0.995870859 | 2.32E-09 | 1604.325 | 765 |
| ENSMUSG00000018999 | Slc35b4 | -1.008654529 | 8.59E-10 | 5598.35 | 2647.85 |
| ENSMUSG00000038072 | Galnt11 | -1.010567151 | 1.41E-09 | 1338.325 | 630.875 |
| ENSMUSG00000042104 | Uggt2 | -1.013830684 | 0.003931886 | 33.45 | 15.625 |
| ENSMUSG00000067370 | B3galt4 | -1.019062733 | 6.19E-08 | 1237.2 | 579.925 |
| ENSMUSG00000074004 | B3gnt6 | -1.021857981 | 0.389860796 | 1.2 | 0.525 |

| | | | | | |
|---|---|---|---|---|---|
| ENSMUSG00000005949 | Ctns | -1.032988997 | 4.86E-08 | 719.8 | 334.45 |
| ENSMUSG00000036356 | Csgalnact1 | -1.033615857 | 1.00E-08 | 1224 | 567 |
| ENSMUSG00000049721 | Gal3st1 | -1.0600812 | 0.181835563 | 2.9 | 1.3 |
| ENSMUSG00000051650 | B3gnt2 | -1.075870159 | 0.000287094 | 80.15 | 35.95 |
| ENSMUSG00000079445 | B3gnt7 | -1.108754507 | 0.007217712 | 16.25 | 7.075 |
| ENSMUSG00000029162 | Khk | -1.128244235 | 4.59E-08 | 478.25 | 207.925 |
| ENSMUSG00000022747 | St3gal6 | -1.184345781 | 1.51E-06 | 594.025 | 248.05 |
| ENSMUSG00000054008 | Ndst1 | -1.193953107 | 3.52E-09 | 4709.6 | 1959.025 |
| ENSMUSG00000026110 | Mgat4a | -1.216928961 | 1.05E-10 | 2695.475 | 1102.05 |
| ENSMUSG00000031749 | St3gal2 | -1.245478396 | 1.78E-09 | 5738.35 | 2305.475 |
| ENSMUSG00000057286 | St6galnac2 | -1.274465901 | 0.000232523 | 69.35 | 27.1 |
| ENSMUSG00000035273 | Hpse | -1.316288683 | 8.73E-06 | 159.975 | 61.075 |
| ENSMUSG00000052544 | St6galnac3 | -1.367806898 | 0.000169799 | 113.6 | 41.675 |
| ENSMUSG00000029431 | B3gnt4 | -1.38027386 | 0.044420378 | 3.675 | 1.3 |
| ENSMUSG00000040710 | St8sia4 | -1.390427185 | 3.49E-10 | 13092.875 | 4756.225 |
| ENSMUSG00000046152 | Fut10 | -1.403272493 | 0.006796863 | 22.275 | 7.925 |
| ENSMUSG00000030711 | Sult1a1 | -1.424667852 | 0.004323491 | 11.6 | 4.1 |
| ENSMUSG00000043857 | Mgat5b | -1.543872547 | 0.305106457 | 0.225 | 0 |
| ENSMUSG00000008461 | Fut1 | -1.544902221 | 0.305640727 | 0.25 | 0 |
| ENSMUSG00000044499 | Hs3st5 | -1.545031112 | 0.305708439 | 0.225 | 0 |
| ENSMUSG00000038296 | Galnt18 | -1.551039025 | 0.309091205 | 0.275 | 0 |
| ENSMUSG00000089704 | Galnt2 | -1.64354506 | 0.022296489 | 6.1 | 1.775 |
| ENSMUSG00000013418 | B4galnt2 | -1.658945442 | 0.019554466 | 103.975 | 31.025 |
| ENSMUSG00000021504 | B4galt7 | -1.688984072 | 3.23E-13 | 3302.325 | 973.95 |
| ENSMUSG00000030657 | Xylt1 | -1.76592356 | 6.34E-08 | 951.175 | 266.3 |
| ENSMUSG00000013846 | St3gal1 | -1.95789188 | 2.98E-11 | 3970.925 | 972.55 |
| ENSMUSG00000005043 | Sgsh | -1.968098375 | 5.59E-14 | 1609.375 | 390.675 |
| ENSMUSG00000021065 | Fut8 | -2.046750492 | 1.59E-14 | 2908.125 | 668.975 |
| ENSMUSG00000026080 | Chst10 | -2.080913788 | 6.64E-13 | 1400.85 | 314.825 |
| ENSMUSG00000020520 | Galnt10 | -2.086089182 | 4.32E-15 | 10273.925 | 2301.175 |
| ENSMUSG00000033849 | B3galt2 | -2.215836996 | 0.000111842 | 34.15 | 6.9 |
| ENSMUSG00000096914 | Galntl6 | -2.272886439 | 0.186001165 | 0.475 | 0 |
| ENSMUSG00000055373 | Fut9 | -2.278974981 | 0.157943352 | 0.5 | 0 |
| ENSMUSG00000016918 | Sulf1 | -2.289353956 | 0.000977507 | 9.65 | 1.75 |
| ENSMUSG00000042082 | Arsb | -2.370480421 | 2.85E-11 | 4616.95 | 851.3 |
| ENSMUSG00000055978 | Fut2 | -2.408137402 | 2.15E-05 | 51.475 | 9.125 |
| ENSMUSG00000022885 | St6gal1 | -2.427756316 | 2.92E-16 | 13120.525 | 2317.8 |
| ENSMUSG00000030283 | St8sia1 | -2.704737987 | 2.07E-13 | 4898.25 | 715.4 |
| ENSMUSG00000047759 | Hs3st3a1 | -3.406497608 | 0.061697342 | 1.225 | 0 |
| ENSMUSG00000031910 | Has3 | -3.451365366 | 1.98E-14 | 1257.95 | 109.25 |
| ENSMUSG00000039037 | St6galnac5 | -3.657315946 | 2.41E-07 | 28.45 | 2.025 |
| ENSMUSG00000026828 | Galnt5 | -3.660937254 | 0.035203225 | 1.575 | 0 |
| ENSMUSG00000074892 | B3galt5 | -3.738431634 | 5.25E-08 | 69.575 | 4.825 |

Supplementary Table 2. Properties of Siglec-1 counter-receptors expressed by activated Tregs

| **Uniprot Entry** | **Gene Name** | **Protein Name** | **Function** | **Adhesion / Signaling** |
|---|---|---|---|---|
| P01849 | Tcra | T-cell receptor alpha chain C region | | |
| Q64253 | Ly6e Ly67 | Lymphocyte antigen 6E | Involved in T-cell development. | |
| P15702 | Spn | Leukosialin; CD43 | Predominant cell surface sialoprotein of leukocytes which regulates multiple T-cell functions, including T-cell activation, proliferation, differentiation, trafficking and migration. Acts as a T-cell counter-receptor for Siglec 1. | Adhesion and Signaling |
| P01831 | Thy1 Thy-1 | Thy-1 membrane glycoprotein; CD90 | May play a role in cell-cell or cell-ligand interactions. | |
| P15379 | Cd44 Ly-24 | CD44 | Cell-surface receptor that plays a role in cell-cell interactions, cell adhesion and migration, helping them to sense and respond to changes in the tissue microenvironment. Engages, through its ectodomain, extracellular matrix components. | Adhesion and Signaling |
| Q08481 | Pecam1 Pecam Pecam-1 | Platelet endothelial cell adhesion molecule; CD31 | Cell adhesion molecule required for leukocyte migration. Promotes macrophage-mediated phagocytosis of apoptotic leukocytes. | Adhesion |
| Q9Z0T9 | Itgb6 | Integrin beta-6 | Mediates R-G-D-dependent release of transforming growth factor beta-1 (TGF-beta-1) from regulatory Latency-associated peptide (LAP), thereby playing a key role in TGF-beta-1 activation | Adhesion and Signaling |
| B2RXS4 | Plxnb2 | Plexin-B2 | Cell surface receptor for SEMA4C, SEMA4D and SEMA4G that plays an important role in cell-cell signaling. | Signaling |
| Q61490 | Alcam | CD166; Activated leukocyte cell adhesion molecule | Cell adhesion molecule that promotes T-cell activation and proliferation. Contributes to the formation and maturation of the immunological synapse. | Adhesion |
| P24063 | Itgal Lfa-1 Ly-15 | Integrin alpha-L; CD11a | Involved in leukocyte adhesion and transmigration of leukocytes including T-cells. | Adhesion |
| Q9Z0M6 | Adgre5 Cd97 | Adhesion G protein-coupled receptor E5; CD97 | Receptor potentially involved in both adhesion and signaling processes early after leukocyte activation. Plays an essential role in leukocyte migration. | Adhesion and Signaling |
| O54901 | Cd200 Mox2 | OX-2 membrane glycoprotein; CD200 | Costimulates T-cell proliferation. May regulate myeloid | Adhesion |
| Q8BTY2 | Slc4a7 Nbc3 | Sodium bicarbonate cotransporter 3 | Electroneutral sodium- and bicarbonate-dependent cotransporter | |
| Q9QUN7 | Tlr2 | Toll-like receptor 2; CD282 | Forms activation clusters composed of several receptors depending on the ligand, these clusters trigger signaling from the cell surface and subsequently are targeted to the Golgi in a lipid-raft dependent pathway. | Signaling |
| O70309 | Itgb5 | Integrin beta-5 | Receptor for fibronectin. | Signaling |
| P18572 | Bsg | Basigin; CD147 | Plays an important role in targeting the monocarboxylate transporters SLC16A1, SLC16A3, SLC16A8, SLC16A11 and SLC16A12 to the plasma membrane. | Adhesion |
| O35235 | Tnfsf11 Opgl Rankl Trance | Tumor necrosis factor ligand superfamily member 11 | Cytokine that binds to TNFRSF11B/OPG and to TNFRSF11A/RANK. Augments the ability of dendritic cells to stimulate naive T-cell proliferation. May be an important regulator of interactions between T-cells and dendritic cells and may play a role in the regulation of the T-cell-dependent immune response. | Signaling |
| O54890 | Itgb3 | Integrin beta-3; CD61 | Receptor for cytotactin, fibronectin, laminin, matrix metalloproteinase-2, osteopontin, osteomodulin, prothrombin, thrombospondin, vitronectin and von Willebrand factor. | Adhesion and Signaling |
| P01590 | Il2ra Il2r | Interleukin-2 receptor subunit alpha; CD25 | Receptor for interleukin-2. The receptor is involved in the regulation of immune tolerance by controlling regulatory T cells activity. | Signaling |
| P01899 | H2-D1 | H-2 class I histocompatibility antigen, D-B alpha chain (H-2D(B)) | Involved in the presentation of foreign antigens to the immune system. | Signaling |
| P01897 | H2-L | H-2 class I histocompatibility antigen, L-D alpha chain | Involved in the presentation of foreign antigens to the immune system. | |
| P04925 | Prnp Prn-p Prp | Major prion protein; CD230 | May be required for neuronal myelin sheath maintenance. May promote myelin homeostasis through acting as an agonist for ADGRG6 receptor. | Signaling |

| | | | | |
|---|---|---|---|---|
| P06800 | Ptprc Ly-5 | Receptor-type tyrosine-protein phosphatase C; CD45 | Protein tyrosine-protein phosphatase required for T-cell activation through the antigen receptor. | Signaling |
| P09055 | Itgb1 | Integrin beta-1; CD29 | Receptors for collagen. | Adhesion and Signaling |
| P10852 | Slc3a2 Mdu1 | 4F2 cell-surface antigen heavy chain; CD98 | Component of several heterodimeric amino acid transporter complexes. When associated with LAPTM4B, the heterodimer formed by SLC3A2 and SLC7A5 is recruited to lysosomes to promote leucine uptake into these organelles, and thereby mediates mTORC1 activation (By similarity). | |
| P11835 | Itgb2 | Integrin beta-2; CD18 | Receptor for ICAM1, ICAM2, ICAM3 and ICAM4. Involved in leukocyte adhesion and transmigration of leukocytes including T-cells (By similarity) | Adhesion |
| P13379 | Cd5 Ly-1 | T-cell surface glycoprotein CD5 | May act as a receptor in regulating T-cell proliferation. | |
| P13597 | Icam1 Icam-1 | Intercellular adhesion molecule 1; CD54 | ICAM proteins are ligands for the leukocyte adhesion protein LFA-1 (integrin alpha-L/beta-2). | Adhesion |
| P14094 | Atp1b1 Atp4b | Sodium/potassium-transporting ATPase subunit beta-1 | This is the non-catalytic component of the active enzyme, which catalyzes the hydrolysis of ATP coupled with the exchange of Na(+) and K(+) ions across the plasma membrane. | |
| P18181 | Cd48 Bcm-1 | CD48 | Ligand for CD2. Might facilitate interaction between activated lymphocytes. Probably involved in regulating T-cell activation. | Signaling |
| P21995 | Emb Gp70 | Embigin | Plays a role in targeting the monocarboxylate transporters SLC16A1 and SLC16A7 to the cell membrane (By similarity). | Adhesion |
| P26011 | Itgb7 | Integrin beta-7 | Involved in adhesive interactions of leukocytes. | Adhesion and Signaling |
| P37217 | Cd69 | CD69 | Involved in lymphocyte proliferation | |
| P55772 | Entpd1 Cd39 | Ectonucleoside triphosphate diphosphohydrolase 1; CD39 | Could hydrolyze ATP and other nucleotides to regulate purinergic neurotransmission. | |
| P56677 | St14 Prss14 | Suppressor of tumorigenicity 14 protein homolog | Degrades extracellular matrix. | |
| P57716 | Ncstn | Nicastrin | Essential subunit of the gamma-secretase complex, an endoprotease complex that catalyzes the intramembrane cleavage of integral membrane proteins such as Notch receptors and APP (amyloid-beta precursor protein). | Signaling |
| P97370 | Atp1b3 | Sodium/potassium-transporting ATPase subunit beta-3; CD298 | Non-catalytic component of the active enzyme, which catalyzes the hydrolysis of ATP coupled with the exchange of Na(+) and K(+) ions across the plasma membrane. | |
| Q00609 | Cd80; B7 | CD80 | Involved in the costimulatory signal essential for T lymphocytes activation. | |
| Q02242 | Pdcd1 Pd1 | Programmed cell death protein 1; CD279 | Inhibitory receptor on antigen activated T-cells that plays a critical role in induction and maintenance of immune tolerance to self. | Signaling |
| Q08857 | Cd36 | Platelet glycoprotein 4; CD36 | Multifunctional glycoprotein that acts as receptor for a broad range of ligands. They are generally multivalent and can therefore engage multiple receptors simultaneously, the resulting formation of CD36 clusters initiates signal transduction and internalization of receptor-ligand complexes. | Signaling |
| Q5FWI3 | Cemip2 Kiaa1412 Tmem2 | Cell surface hyaluronidase | Cell surface hyaluronidase that mediates the initial cleavage of extracellular high-molecular-weight hyaluronan into intermediate-size hyaluronan. | Signaling |
| Q9WVS0 | Icos Ailim | Inducible T-cell costimulatory; CD278 | Enhances all basic T-cell responses to a foreign antigen, namely proliferation, secretion of lymphokines, up-regulation of molecules that mediate cell-cell interaction. Prevents the apoptosis of pre-activated T-cells. | |
| Q60677 | Itgae | Integrin alpha-E | Receptor for E-cadherin. | Adhesion |
| Q61098 | Il18r1 | Interleukin-18 receptor 1; CD218a | Within the IL18 receptor complex, responsible for the binding of the proinflammatory cytokine IL18. | |
| Q61503 | Nt5e Nt5 Nte | 5'-nucleotidase; CD73 | Hydrolyzes extracellular nucleotides into membrane permeable nucleosides. | |

| | | | | |
|---|---|---|---|---|
| Q64735 | Cr1l Crry Cry | Complement component receptor 1-like protein | Acts as a cofactor for complement factor I. Also acts as a costimulatory factor for T-cells which favors IL-4 secretion. | |
| Q80UL9 | Jaml Amica1 Gm638 | Junctional adhesion molecule-like | Transmembrane protein of the plasma membrane of leukocytes that control their migration and activation. | Adhesion and Signaling |
| Q9EP73 | Cd274 B7h1 Pdcd1l1 Pdcd1lg1 Pdl1 | Programmed cell death 1 ligand 1; CD274 | Plays a critical role in induction and maintenance of immune tolerance to self. Modulates the activation threshold of T-cells and limits T-cell effector response. | |
| Q9QUM4 | Slamf1 Slam | Signaling lymphocytic activation molecule; CD150 | Self-ligand receptor of the signaling lymphocytic activation molecule (SLAM) family. SLAM receptors triggered by homo- or heterotypic cell-cell interactions are modulating the activation and differentiation of a wide variety of immune cells and thus are involved in the regulation and interconnection of both innate and adaptive immune response. | Signaling |
| Q9Z127 | Slc7a5 Lat1 | Large neutral amino acids transporter small subunit 1 | The heterodimer with SLC3A2 functions as sodium-independent, high-affinity transporter that mediates uptake of large neutral amino acids. When associated with LAPTM4B, the heterodimer formed by SLC3A2 and SLC7A5 is recruited to lysosomes to promote leucine uptake into these organelles, and thereby mediates mTORC1 activation. | |
